# Insulin stimulation induces dynamic interaction between 14-3-3ε and the C-terminal tail domain of Kinesin-2 subunit KIF3B in vitro

**DOI:** 10.1101/2025.08.20.671056

**Authors:** Diksha Kumari, Anvi Tangri, Mukul Girotra, Krishanu Ray

**Affiliations:** Tata Institute of Fundamental Research, Mumbai 400005, India; Birla Institute of Technology and Sciences, Pilani (BITS-Pilani), Hyderabad, 500078, India

## Abstract

Heterotrimeric kinesin-2 mediates diverse intracellular transport processes, yet the mechanisms governing its cargo specificity and regulation remain poorly understood. In this study, we report the regulation of interaction between the kinesin-2 subunit KIF3B and a candidate adaptor protein, 14-3-3ε. Using split-APEX2 proximity labelling and fluorescence lifetime imaging microscopy (FLIM) assays in HEK293T cells, we demonstrate that 14-3-3ε interacts with KIF3B within discrete subcellular compartments. This interaction is significantly enhanced upon insulin stimulation, indicating regulation by signalling pathways. Time-resolved FLIM analysis further reveals that the site of interaction is dynamic, suggesting spatial and temporal control of motor–adaptor association. Finally, the interactions are mapped to two serine residues in the C-terminal tail region of KIF3B that are highly conserved across species. Together, these findings uncover a conditional and reversible interaction between KIF3B and 14-3-3ε, providing insight into how kinesin-2 may achieve context-specific cargo engagement and transport within cells.

**Highlights of the study:**

- 14-3-3ε binds to KIF3B subunit of kinesin-2 in restricted subcellular compartments
- Insulin stimulation triggers the association between 14-3-3ε and KIF3B
- Ser631 and Ser640 on KIF3B are the potential target sites of 14-3-3ε binding

## Introduction

The interaction between a motor protein and its cargo must be transient to enable selective and directional transport in response to any physiological stimuli. This principle is well documented for kinesin motors, which execute active intracellular transport of a wide range of cargo, including soluble proteins, cellular organelles, secretory vesicles, and RNA–protein complexes ^1^. Kinesins engage their cargo in a dynamic and spatiotemporally regulated manner, binding and releasing at defined subcellular sites in response to signalling cues. Neuronal activity, for example, has been shown to regulate the transient interaction between kinesin⍰2 and the soluble enzyme choline acetyltransferase (ChAT) within axons, resulting in increased ChAT flux toward the axon terminus through a combination of active and passive mechanisms^2,3^. A similar regulated interaction occurs between kinesin⍰2 and N⍰cadherin. Under chronic neuronal inactivity, kinesin-2 subunit KIF3A is phosphorylated by PKA and CaMKII, leading to N⍰cadherin recruitment and subsequent transport toward dendritic spines in mouse hippocampal neurons ^4^. Such regulated motor–cargo associations are frequently controlled by extracellular and intracellular signalling pathways that modulate kinesin phosphorylation states (reviewed in ^5^). Biochemical studies further suggested that insulin stimulation could activate Rab4 in adipocytes, facilitating kinesin⍰2 recruitment and translocation of Glut4 vesicles to cell periphery through an as⍰yet⍰undefined mechanism^6^. Identifying these transient associations require time⍰resolved protein–protein interaction assays in living cells or differentiated tissues, rather than conventional *in vitro* pulldown approaches, which capture only static snapshots and often fail to detect weak or short⍰lived interactions occurring *in vivo*.

14⍰3⍰3 proteins constitute a highly conserved family of small homo⍰ and heterodimeric scaffolding proteins (∼30 kDa) that bind a broad spectrum of target proteins through a characteristic amphipathic groove ^7^. They participate in diverse signalling pathways, including apoptosis, autophagy, metabolism, cell⍰cycle control, and intracellular transport, by recognizing consensus phospho⍰motifs and acting as intermediaries that couple signalling events to downstream cellular responses^8^. For instance, insulin stimulation promotes phosphorylation⍰dependent binding of 14⍰3⍰3 proteins to IRS1 and IRS2, establishing a negative feedback loop that modulates insulin signalling ^9^. 14⍰3⍰3 proteins also interact with multiple kinesin motors and their cargoes, thereby influencing motor–cargo association and kinesin function^10–15^. In *Drosophila* oocytes, the 14⍰3⍰3ζ and 14⍰3⍰3ε isoforms bind the kinesin⍰3 orthologue KHC73 and the dynein adaptor NudE, respectively, where their dimerization contributes to mitotic spindle organization^10^. In neurons, 14⍰3⍰3 binding to MARK2⍰phosphorylated KIF13A prevents its association with transferrin receptors in axons, ensuring polarized transport toward dendrites ^13^.

Signalling⍰dependent 14⍰3⍰3 associations are further implicated in vesicular trafficking, including the insulin⍰stimulated movement of GLUT4⍰containing vesicles toward the plasma membrane^16,17^. These observations collectively suggest that 14⍰3⍰3 proteins may recruit kinesin⍰family motors to specific vesicle populations in response to insulin signalling. Furthermore, biophysical analyses using time⍰resolved FRET ^18,19^ and related approaches have revealed that 14⍰3⍰3 proteins interact dynamically with phosphorylated target peptides (reviewed in ^20^). Therefore, these proteins provide an ideal candidate for studying dynamic interaction between kinesin motor and its cargo.

In the present study, we examined the dynamics of the interaction between the mammalian kinesin⍰2 subunit KIF3B and 14⍰3⍰3ε in cultured cells to better understand how kinesin⍰2 associates with its cargoes or adaptor proteins. Using time⍰resolved FLIM⍰FRET and enzyme⍰activity⍰based split⍰APEX2 assays, we detected a weak interaction between 14⍰3⍰3ε and KIF3B within specific subcellular compartments of HEK293T cells, which was markedly enhanced upon insulin stimulation. We further identified two candidate 14⍰3⍰3⍰binding sites on KIF3B, centred around Ser631 and Ser640 in the predicted tail domain, that contribute to the insulin⍰dependent interaction between 14⍰3⍰3ε and KIF3B. Together, these findings reveal a signalling⍰dependent, reversible interaction between KIF3B and 14⍰3⍰3ε, establishing a regulatory mechanism through which kinesin⍰2 may selectively engage its cargo or adaptor proteins in response to metabolic cues.

## Results

### 14-3-3 interacts with the tail domain of kinesin-2 subunit KIF3B

To identify whether 14-3-3 proteins could interact with kinesin-2, we used an online prediction tool, <u>14-3-3-Pred</u>, to check for the presence of potential 14-3-3 binding sites on all three kinesin-2 subunits ^21^. We identified two highly conserved potential binding sites in the predicted C-terminal tail domains of KIF3B and its orthologues (Figure 1A), one site on the motor domain of KIF3A and five sites on its accessory protein, called kinesin-associated protein (KAP) **(Table1)**. Kinesin-family motors involved in long-range intracellular transport, bind to their adaptor/cargo complex through their tail domain, while the motor domain mediates its interaction with microtubules ^22–24^. Hence, for this study, we focused on studying the interaction of 14-3-3ε with KIF3B subunit of kinesin-2. Sequence alignment of predicted 14-3-3ε binding sites on KIF3B revealed that the two serine residues, Ser631 and Ser640, which lie within the first 51 amino acid (1/3^rd^) segment of the tail domain of human KIF3B, are highly conserved across different species including *Drosophila* KLP68D **(Figure 1A)**.

**Table 1:** Potential 14-3-3 binding sites on kinesin-2 subunits.

| **Kinesin-2 subunit** | **Phosphorylatable residue** | **Potential binding motif** | **ANN** | **PSSM** | **SVM** | **Consensus** |
|---|---|---|---|---|---|---|
| MmKIF3A | Thr200 | DMDRIM[T]LGHK | 0.803 | 1.079 | 0.918 | 0.933 |
| MmKIF3B | Ser631 | MMKRPV[S]AVGY | 0.814 | 1.031 | 0.695 | 0.847 |
|  | Ser640 | GYKRPL[S]QHAR | 0.730 | 0.889 | 0.486 | 0.702 |
| MmKAP3 | Ser60 | KIIRLK[S]LNAN | 0.845 | 1.149 | 1.110 | 1.035 |
|  | Ser102 | LQNRRD[S]LSGK | 0.688 | 0.814 | 0.493 | 0.665 |
|  | Ser418 | MDDRFK[S]MFAY | 0.749 | 1.073 | 0.628 | 0.817 |
|  | Ser491 | KMIRNI[S]QHDG | 0.720 | 0.968 | 0.271 | 0.653 |
|  | Thr776 | ILGRPA[T]AYGF | 0.658 | 0.847 | 0.460 | 0.655 |

*ANN - Artificial Neural Network (cut-off = 0.55)*
*PSSM - Position-Specific Scoring Matrix (cut-off = 0.80)*
*SVM - Support Vector Machine (cut-off = 0.25)*
*Consensus - Average of the scores provided by the three methods (cut-off = 0.50)*

**Figure 1.**
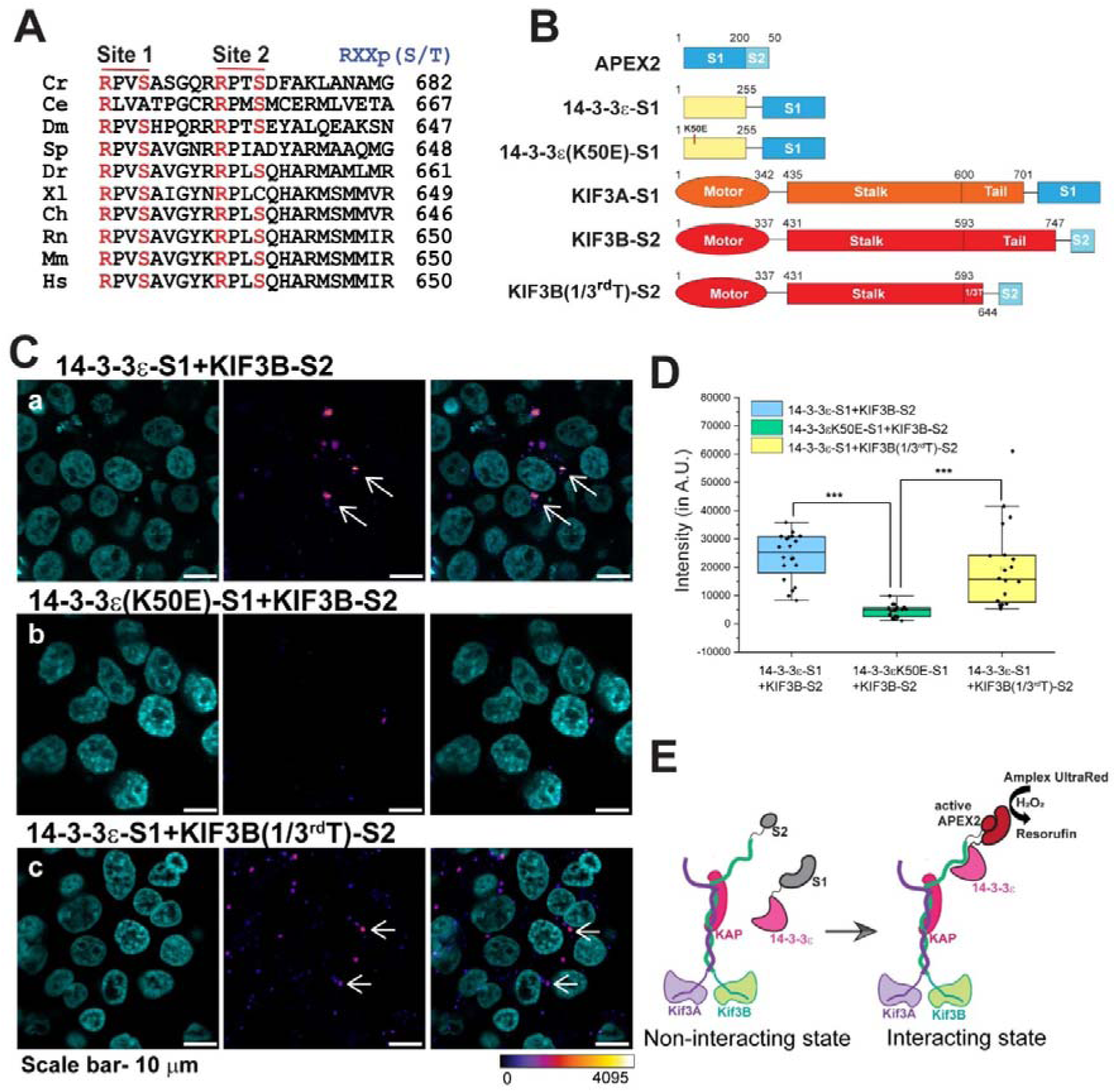
Split-APEX2 assay shows 14-3-3ε interacts with KIF3B in a specific sub-cellular region. **A.** Alignment of potential 14-3-3-binding sites Ser631 (Site 1) and Ser640 (Site 2) on the KIF3B tail domain and its orthologues in different species. The conserved serine and arginine residues in the predicted sites are marked in red. Both the sites lie within the first 51 amino acid fragment of the C-terminal tail domain of KIF3B, which is indicated as 1/3^rd^ Tail in panel B. ***Glossary:*** *Cr-Chlamydomonas reinhardtii, Ce*-*Caenorhabditis elegans, Dm-Drosophila melanogaster, Sp-Strongylocentrotus purpuratus, Dr-Danio rerio, Xl-Xenopus laevis, Ch-Capra hircus, Rn-Rattus norvegicus, Mm-Mus musculus, Hs-Homo sapiens* **B.** Schematic representation of constructs cloned in the pCMV vector and used for the split-APEX2 assay. **C.** Images show the reconstitution of APEX2 activity in HEK293T cells transfected with **a)** 14-3-3ε-S1 and KIF3B-S2 (test), **b)** 14-3-3ε(K50E)-S1 and KIF3B-S2 (negative control), and **c)** 14-3-3ε-S1 and KIF3B(1/3^rd^T)-S2 (to map the binding site), as mentioned above each panel. The fluorescence intensity of resorufin produced by the oxidation of Amplex UltraRed (AUR) due to APEX2 activity on H_2_O_2_ is shown in false color heat map as per the scale on the bottom. The Hoechst dye (cyan) marked the nuclei. Arrows indicate 14-3-3ε-S1 and KIF3B-S2 interaction sites and scale bars ∼10 µm. **D.** Quantification of the resorufin signals in different backgrounds shown in panel C (as described in the color code in the legend) on a relative scale indicated on the y-axis. Each point represents an ROI drawn for 10 nuclei. The pair-wise significance of difference was estimated using Mann-Whitney U test, *** indicates p value < 0.001. The central line shows the median. The whiskers show the values extending to 1.5X the interquartile range from the Q1 or Q3 boundaries. N=3, n=20 ROIs (200 cells).

To test the interaction between KIF3B and 14-3-3ε we first used the split-APEX2 assay. Ascorbate Peroxidase (APEX) is a fast-acting enzyme, which produces short-lived highly reactive free radicals in the presence of H_2_O_2_ ^25,26^. Reactive oxygen species can convert biotin-phenol into biotin-phenoxy radicals that can attach to electron-rich amino acid side chains, such as lysine residues, of the nearby proteins, thereby biotinylating them^27^. APEX2 is an engineered variant with improved catalytic efficiency and sensitivity^28^. A split version of APEX2 was subsequently developed, in which the enzyme is divided into two individually inactive fragments that can reconstitute enzymatic activity when brought into proximity through interacting fusion partners^29^. The reconstituted peroxidase activity can be detected using fluorogenic substrates, such as Amplex UltraRed, which is oxidized to a red fluorescent Resorufin in the presence of reactive free radicals ^29^. Thus, split-APEX2 assay involving two separate proteins tagged with inactive enzyme fragments, referred to as S1 and S2 in this study, could confirm the association between the two proteins in cellular contexts.

To utilize this assay for testing potential interaction between 14-3-3ε and KIF3B, we tagged the 200 aa N-terminal APEX2 fragment (S1) at the C-terminal end of 14-3-3ε and the 50 aa C-terminal APEX2 fragment (S2) at the C-terminal end of KIF3B **(Figure 1B)**. We reasoned that association between 14-3-3ε and the KIF3B is likely to bring in the APEX2 fragments close together and reconstitute the peroxidase activity which will be reported by the conversion of Amplex UltraRed to insoluble Resorufin precipitate at the sites of interactions. We also reasoned that APEX2 being highly sensitive would be able to capture even the weak and transient interactions that are often not captured by standard pull-down assays^29–31^. The assay was first validated using full length APEX2 enzyme tagged to mitochondria localization signal (mito-V5-APEX2) ^27^, which produced a strong red fluorescence signal distributed throughout the cytoplasm, characteristic of mitochondria staining observed in previous studies^25,29,32^ (**Figure S1A**). Furthermore, using KIF3A-S1 and KIF3B-S2 pair as positive control of split-APEX2 assay also produced a strong red fluorescence signal in the cytoplasm of the co-transfected cells **(Figure S1A).** The negative control pair of untagged S1 and KIF3B-S2 produced no detectable signal **(Figure S1A, C)**. Together, this data indicated that the split-APEX2 assay could be used to study the interaction of two different proteins in tissue cultured cells. It also indicated that the S1 and S2 fragments alone cannot bind each other to reconstitute the enzyme activity in the absence of the tags.

Next, we tested split-APEX2 reconstitution using the 14-3-3s-S1 and KIF3B-S2. The cells transfected with the 14-3-3ε-S1 and KIF3B-S2 pair produced a modest fluorescence signal localized to a small cytoplasmic compartment **(arrows, Figure 1 C-a, Figure S1B)**, suggesting that 14-3-3ε and KIF3B could associate with each other in tissue-cultured cells within a select compartment. In comparison, the mutant 14-3-3ε(K50E)-S1, carrying a point mutation converting Lys50 to Glu in the ligand binding amphipathic groove that disrupts binding to phosphorylated Ser/Thr residues, produced no resorufin signal in the presence of KIF3B-S2 **(Figure 1 C-b).** The KIF3B(1/3^rd^T)-S2 construct expressing KIF3B with only the proximate 51 amino acid of the C-terminal tail domain (Figure 1B), containing the predicted S631 and S640 residues, could also produce the signal when co-expressed with 14-3-3ε-S1 **(Figure 1 C-c)**. The signal quantification revealed that both wild type (14-3-3□-S1 and KIF3B-S2) and the tail-truncated KIF3B (14-3-3□-S1 and KIF3B(1/3^rd^T)-S2) pairs produced significantly higher signals than the negative control pair (14-3-3ε(K50E)-S1 and KIF3B-S2), respectively **(Figure 1D).** Together, the split-APEX2 assay suggested a potential interaction between 14-3-3ε and kinesin-2 in cellular conditions (Figure 1E). However, the possibility of a fixation induced artefact could not be ruled out.

### Fluorescence lifetime Imaging Microscopy (FLIM) revealed 14-3-3ε interacts with KIF3B in distributed subcellular compartments

Therefore, we utilized the Fluorescence Lifetime Imaging Microscopy (FLIM) assay to assess the interaction between the KIF3B tagged with mTurquoise2 (mTq2) and 14-3-3ε tagged with mCitrine (mCit) in live HEK293T cells. We used cells transfected with the KIF3B-mTq2 alone to estimate the base level for comparison. First, we tested the conditions with a positive control of KIF3AmCit and KIF3BmTq2 pair. These two proteins are known to form an obligate heterodimer. We used phasor plots, a fit-free representation of the fluorescence lifetime data where each point represents the lifetime of a pixel mapped to the G (cosine) and S (sine) components derived after Fourier-transformation of the decay curves^33,34^ to estimate the gross change in lifetime at each pixel. Points lying on the universal semicircle of a phasor plot indicate single component fluorescence decay, while those lying inside the universal semicircle represent multi-component lifetime decay^33^. Thus, it provides an intuitive estimate of the global distribution without curve fitting.

In **FiguresS2A-c** and **S2B-c**, the phasor distribution was predominantly localized along the universal semicircle, consistent with an approximately mono-exponential fluorescence decay. The cluster centred towards a lower lifetime in the KIF3AmCit and KIF3BmTq2 background (**FigureS2B-c**) as compared to KIF3BmTq2 alone (**Figure S2A-c**). Similarly, the frequency histogram analysis of pixel-wise lifetime distribution in the cytoplasmic region of the cells expressing both the proteins indicated a clear shift towards lower lifetime **(Figure S2C)**. The mean lifetime decreased significantly from 4031.0 ± 73 ps to 3713.5 ± 97 ps **(Figure S2D)**, suggesting an increased association between the tagged proteins. This reduction in donor fluorescence lifetime was consistent with the F⍰rster’s resonance energy transfer (FRET) occurring between mTq2 and mCit ^35^, indicating a relative proximity between KIF3B and KIF3A in living cells. These results motivated us to further investigate protein–protein interactions involving KIF3BmTq2 by quantifying fluorescence lifetime changes in the presence of 14-3-3εmCit.

The average KIF3BmTq2 lifetime values were not significantly reduced in the presence of 14-3-3εmCit, as compared to 14-3-3ε(K50E)mCit **(Figure 2A-D).** However, the FLIM data revealed a heterogeneous distribution of lifetime within the cytoplasm, with some localized regions of lower lifetime specifically in the presence of 14-3-3εmCit, suggesting that the association between KIF3B and 14-3-3ε could be restricted to small subcellular compartments **(arrows in Figure 2A c, d)**.

**Figure 2.**
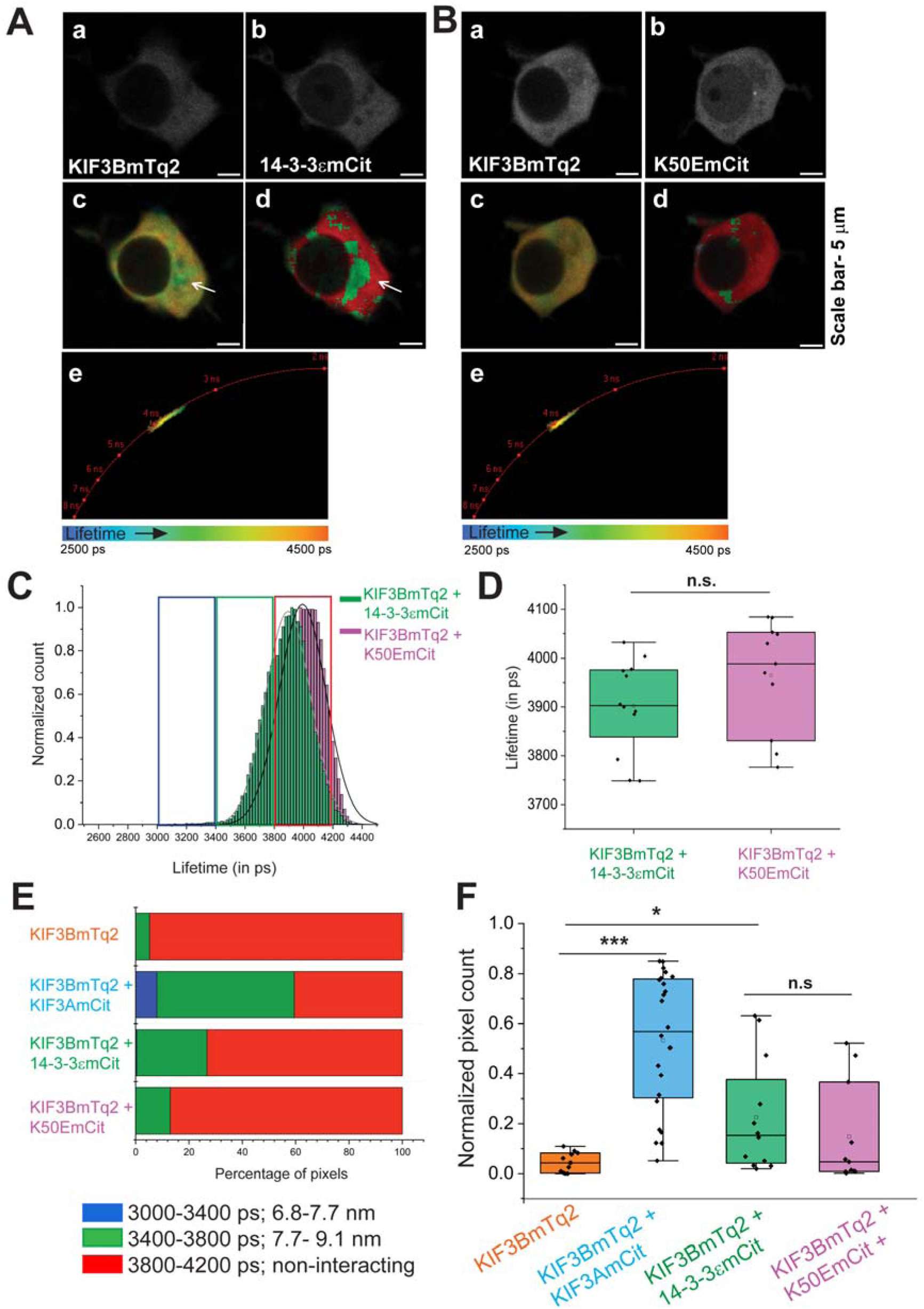
Studying the interaction of KIF3B with 14-3-3ε using FLIM-FRET. **A.** Fluorescence lifetime measurement of KIF3BmTq2 in the presence of 14-3-3εmCit. **(a, b)** Confocal images of KIF3BmTq2 and 14-3-3εmCit, respectively; **(c)** fluorescence lifetime image of KIF3BmTq2; **(d)** false-colour image showing the lifetime of KIF3BmTq2 in discrete value ranges, where red represents lifetime values in the range of 3800–4200 ps and green represents lifetime values in the range of 3400–3800 ps; **(e)** phasor plot shows the lifetime distribution of KIF3BmTq2. Scale bar ∼5 μm. **B.** Fluorescence lifetime measurement of KIF3BmTq2 in the presence of the mutant 14-3-3ε (K50E)mCit. **(a, b)** Confocal images of KIF3BmTq2 and 14-3-3ε (K50E)mCit, respectively; **(c)** fluorescence lifetime image of KIF3BmTq2; **(d)** false-colour image showing the lifetime of KIF3BmTq2 in discrete value ranges as described in A(d); **(e)** phasor plot showing the lifetime distribution of KIF3BmTq2. Scale bar ∼5 μm. **C.** Histogram plots showing the frequency distribution of the fluorescence lifetime of KIF3BmTq2 in the presence of wild-type (green) and the K50E mutant form of 14-3-3εmCit (pink). Red, green and blue boxes demarcate the lifetime values in the range of 3800-4200 ps, 3400-3800 ps and 3000-3400 ps, respectively. **D.** The box and whisker plots depict the lifetime values estimated in different backgrounds as indicated on the x-axis as described in Figure 1D. The statistical significance of difference was estimated using Mann-Whitney U test. n.s. indicates p value > 0.05. The central line shows the median. The whiskers show the values extending to 1.5X the interquartile range from the Q1 or Q3 boundaries. n≥10 cells. **E.** Histogram stack of percentage pixels having fluorescence lifetime of KIF3BmTq2 in the ranges of 3000–3400 ps (blue), 3400–3800 ps (green), and 3800–4200 ps (red), in the presence of different interacting partners as indicated along the y-axis. **F.** Box and whiskers plots showing the fraction of pixels with the fluorescence lifetime of KIF3BmTq2 in the range of 3400–3800 ps (green band) for each of the indicated conditions along the x-axis. The statistical significance of difference was estimated using Mann-Whitney U test. * Indicates p-value < 0.05, *** indicates p-value < 0.001 and n.s. p-value > 0.05.

To estimate the heterogeneous lifetime distribution in different co-transfection conditions, we divided the pixel-wise lifetime values into three broad categories, according to the lifetime distribution. Free KIF3BmTq2 lifetime fell within 3800 - 4200 ps, corresponding to estimated FRET distance greater than 9.1 nm for a single exponential decay **(Figure S2C, Figure 2E)**. We classified it as non-interacting band (red band). For KIF3BmTq2 and KIF3AmCit pair, the distribution was spread from 3000 - 4200 ps **(Figure S2C, Figure 2E)**. We divided this distribution into two bands - green band (between 3400-3800 ps, corresponding to 7.7- 9.1 nm F⍰rster’s radii for a mono-exponential decay), and blue band (between 3000 – 3400 ps, corresponding to 6.8 - 7.7 nm F⍰rster’s radii). The F⍰rster’s radius (R_0_), which is the distance between a fluorescent donor and acceptor at which 50% of the donor’s excited-state energy is transferred to the acceptor through FRET, for the mTq2 and mVenus pair has been estimated at 5.8 nm ^36^, which is the closest the two fluorochrome could get to each other when the two proteins are at physical contact. Further, the FLIM-FRET estimate for KIF3AmCit and KIF3BmTq2 pair was consistent with the previously estimated FRET distances between the KIF3B/KIF3A stalk heterodimer ^37,38^. Hence, we categorised the other two bands as weak (green) and moderately (blue)-interacting bands, respectively to estimate relative interaction strength.

Pixels with fluorescence lifetime in green zone, indicative of weak interaction, were more prominent with KIF3BmTq2 and 14-3-3εmCit than with the negative control, 14-3-3ε(K50E)mCit **(Figure 2 A, B, E and F)**. Though not significant, both the phasor plot and frequency distribution graphs **(Figure 2A-D**) indicated a shift in the mean lifetime of KIF3BmTq2 to 3901.8 ± 96 ps in the presence of the wild type 14-3-3εmCit as compared to 3964.8 ± 113 ps in the presence of 14-3-3ε(K50E)mCit. Similarly, there was a modest increase in the pixel count in green band in the presence of the 14-3-3εmCit as compared to that of 14-3-3ε(K50E)mCit **(Figure 2F)**. Together, the analysis of the FLIM-FRET data suggested a potentially weak interaction between 14-3-3εmCit and KIF3B restricted to a small perinuclear compartment in the cytoplasm. The data, however, did not rule out a possibility of passive colocalization of the two proteins in the subcellular compartment due to overexpression.

### Insulin stimulation enhances the interaction between 14-3-3ε and KIF3B

The weak interactions between 14-3-3εmCit and KIF3BmTq2 within discrete cytoplasmic domain also implied spatiotemporal regulation of their association. A previous study showed that insulin stimulation to cultured adipocytes activates Rab4-GTPase, increasing its association with kinesin-2, leading to an increase in kinesin-2-dependent trafficking of Glut4 vesicles towards the cell periphery ^6^. Insulin stimulation also increased Glut4 trafficking by recruiting 14-3-3ε/θ to a RabGAP-As160 ^17^. Furthermore, insulin stimulation has been shown to promote association of different 14-3-3 isoforms with their substrates, including Insulin Receptor Substrate, IRS-2 ^39^, and the small GTPase-activating protein GARNL1/RalGAPα1^16^, in the context of GLUT4 trafficking in muscle cells, as well as with a diverse range of other proteins ^40^. Therefore, we decided to test whether the interaction between 14-3-3ε and KIF3B could be enhanced by insulin stimulation in HEK293T cells. For this purpose, we treated the HEK293T cells with 17 nM insulin (approximately 100 ng/mL)^6^, and performed the fluorescence lifetime imaging of KIF3BmTq2 in cells expressing both the fluorophores **(Figure 3A)**, before and after the insulin stimulation.

**Figure 3.**
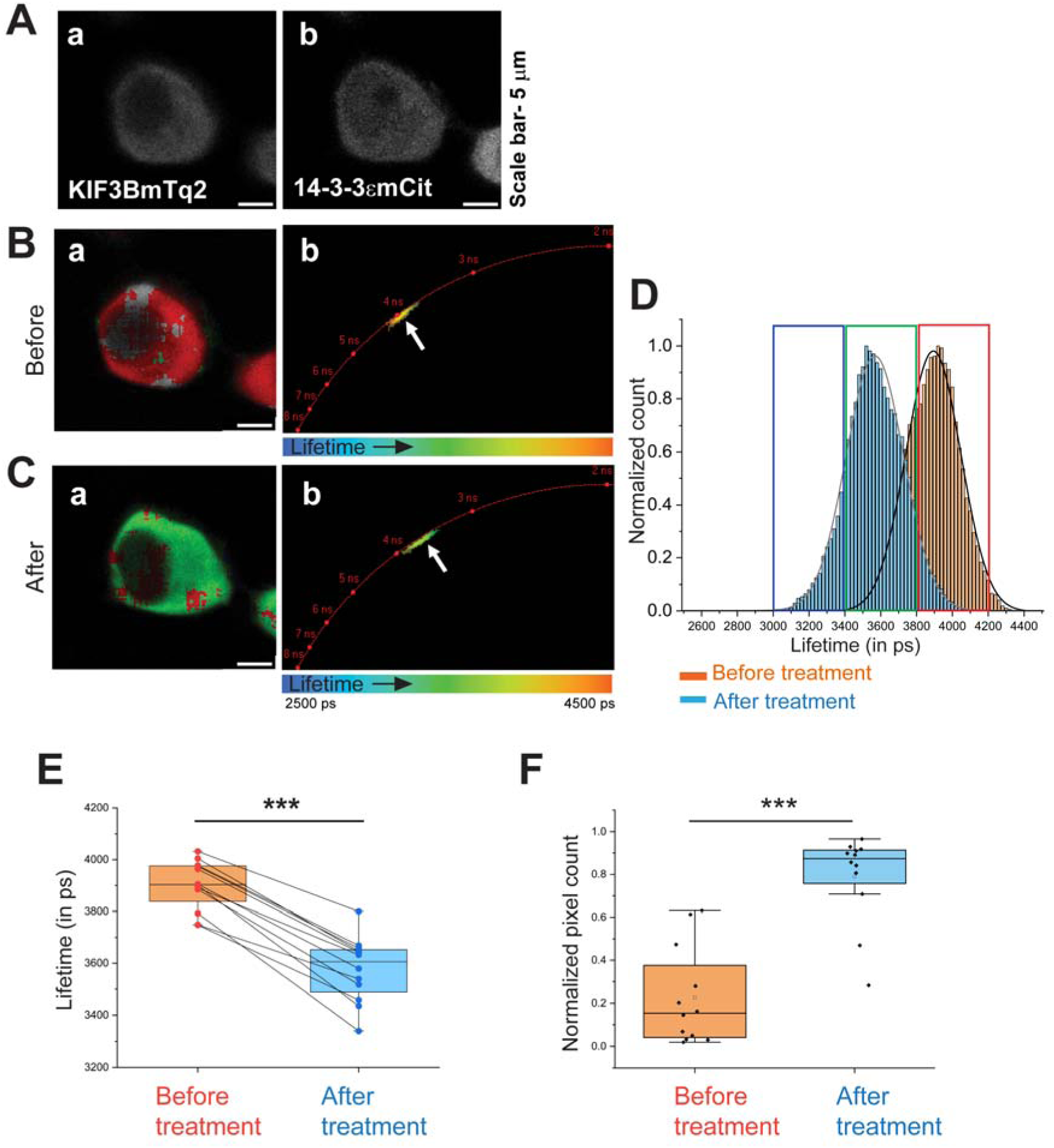
Insulin stimulation increases the interaction between KIF3B and 14-3-3ε. **A.** Confocal images showing the expression of KIF3BmTq2 **(a)** and 14-3-3εmCit **(b)** in a representative HEK293T cell. Scale bar ∼ 5 μm. **B. (a)** False-colour image of the selected cell showing the lifetime of KIF3BmTq2 in discrete value ranges before insulin treatment, where red represents lifetime values in the range of 3800–4200 ps and green represents lifetime values in the range of 3400–3800 ps; **(b)** phasor plot showing the lifetime distribution of KIF3BmTq2. **C. (a)** False-colour image of the same cell showing the lifetime of KIF3BmTq2 in discrete value ranges after insulin treatment (17 nM, for 5 minutes), according to the colour scheme as described in B(a); **(b)** phasor plot showing the lifetime distribution of KIF3BmTq2. **D.** Histogram plot showing the frequency distribution of the fluorescence lifetime of KIF3B-mTq2 before (orange) and after (blue) the insulin treatment. **E.** Box and whiskers plots showing the change in fluorescence lifetime of KIF3BmTq2 in each cell before and after insulin treatment. **F.** Box and whiskers plots showing the fraction of pixels with the fluorescence lifetime of KIF3BmTq2 in the range of 3400–3800 ps, for each cell before and after insulin stimulation. The statistical significance of difference was estimated using paired t-test. *** indicates p-value < 0.001.

Insulin stimulation lowered the lifetime values uniformly throughout the cytoplasm and there was a clear shift of the cluster of dots towards the lower lifetime values in the phasor plot **(Figure 3B, C)**. Additionally, the peak of the pixel-wise lifetime distribution frequency, aggregated across all cells following insulin stimulation, shifted toward the lower values **(Figure 3D).** We also observed a significant decrease in the average lifetime values in the conjugated-cell assay following insulin stimulation **(Figure 3E)**, and a significant increase in the proportion of green band pixels **(Figure 3F).** The mean lifetime value dropped from 3901.8 ± 96 ps to 3576.5 ± 125 ps. To test whether the insulin-dependent reduction in the mean lifetime values was caused due to a specific effect leading to increased association between KIF3BmTq2 and 14-3-3εmCit and rule out the possibility of an increased passive colocalization, we performed the same experiment using KIF3BmTq2 and 14-3-3ε(K50E)mCit. We found no significant change in the distribution of lifetime values before and after the insulin stimulations in the presence of the mutant protein **(Figure S3A, B)**. A similar result was obtained with the blank stimulation control **(Figure S3C, D)**. Altogether, the data suggested that insulin stimulation could increase the active interaction between KIF3B and 14-3-3ε and ruled out the possibility of passive colocalization due to the overexpression after insulin stimulation.

To understand how the 14-3-3ε/KIF3B compartment form after insulin stimulation, we recorded the time-resolved FLIM of KIF3BmTq2 in the presence of 14-3-3εmCit in individual cells over a period of one hour, at 5-minutes intervals in the absence (Untreated) and presence of insulin (with Insulin). We observed that the weakly interacting (green band) compartments were maintained in dynamic fashion with lifetime values oscillating between green and red bands in absence of insulin **(arrows, Figure 4A)**. Following insulin stimulation, there was a significant reduction in the average lifetime in these perinuclear compartments, which was maintained until one hour **(Figure 4B, C).** At the same time, proportion of green pixels increased significantly in the cytoplasm **(Figure 4C)**. Further, it appeared to consolidate the association and elevate the interaction to moderate level (blue band) in a perinuclear compartment in a dynamic fashion **(arrows, Figure 4B)**. Also, the combined proportion of blue and green pixels increased significantly following insulin stimulation (**Figure 4D**). These observations suggested that the insulin treatment could promote and stabilize the interaction between 14-3-3ε and kinesin-2.

**Figure 4.**
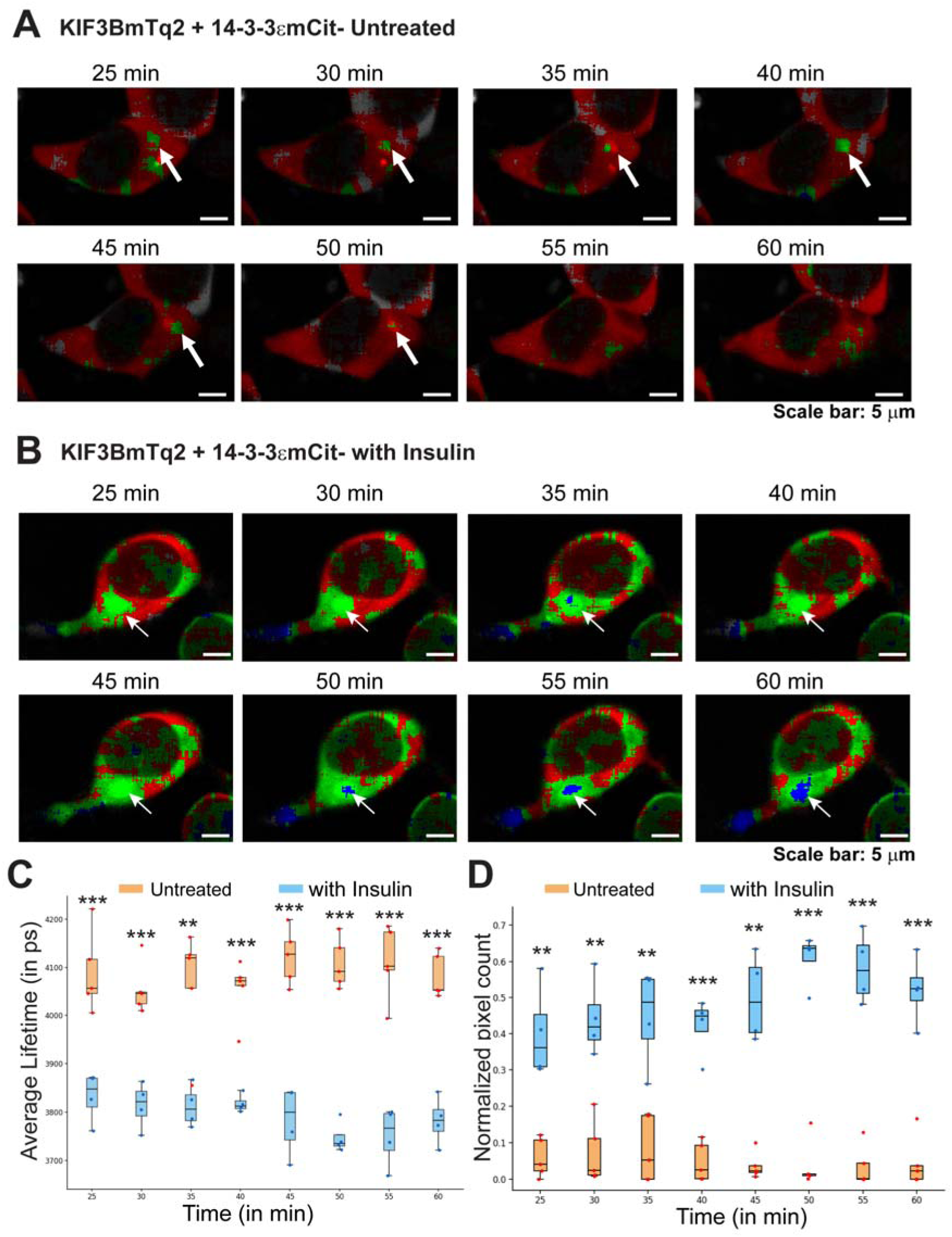
Time course analysis of fluorescence lifetime of KIF3BmTq2 in presence of 14-3-3εmCit in untreated and insulin-treated cells. **A, B.** Time-lapse of fluorescence lifetime map of KIF3BmTq2 in cells coexpressing 14-3-3εmCit in untreated cells **(A)**, and after 17 nm insulin treatment **(B)**. Images are shown in false colour FRET maps as per the colour codes described in Figure 2C, E. Arrows indicate the dynamic interaction compartments within the cell at different time points during the time-course recording. Scale bars ∼ 5 μm. **C.** Box and whisker plots showing average fluorescence lifetime of KIF3BmTq2 in each cell over a period of one hour in an untreated cells (orange) and in cells after 17 nM insulin treatment (blue). The box and whisker plots indicate the lifetime distribution of n=5 cells for untreated and n=4 cells after Insulin treatment. The experiment was performed in five independent sets (N=5) with three showing the effect. **D.** The proportion of pixels with modest FLIM-FRET corresponding to the range of 3000 – 3800 ps in untreated cells (orange) and after 17 nm Insulin treatment (blue). Note that the gross interacting zones expanded by as much as 7 folds after the insulin treatment.

14-3-3 is well known to bind to several of its target proteins that are phosphorylated by certain kinases during cell division ^41,42^. Additionally, the KIF3A/KIF3B/KAP3 complex has also been shown to localize on the mitotic apparatus and facilitate spindle formation and chromosome segregation during cell division ^43^. So far, all these results described were carried out in cells in the presence of FBS and we found noticeable variation in the intensity of the interactions amongst the cells. Therefore, to test whether cell cycle states or serum stimulation could influence the 14-3-3ε and KIF3B association, we studied the interaction in cells after 16 hours of serum starvation, which is likely to arrest the cells in G0 state. Although we observed some compartments of weak interaction **(arrow in Figure S4A-c)**, like the cells grown in presence of FBS, the insulin-dependent enhancement of the interaction between the two proteins was not visible in the cells after serum-starvation **(Figure S4Ac’)**. The proportion of green pixels in serum-starved cells expressing KIFBmTq2 and 14-3-3εmCit remained unchanged after insulin stimulation **(Figure S4B)**. Altogether, these results suggested that insulin stimulation is likely to activate the interaction between KIF3B and 14-3-3ε in the presence of other serum factors and/or in the proliferative state of the cells.

### Two specific serine residues in the KIF3B tail are responsible for the insulin-dependent interaction with 14-3-3ε

To further map the target sites of the interaction, we substituted the S631 and S640 residues of MmKIF3B tail that are predicted to participate in 14-3-3 binding, with alanine in two separate KIF3B(S631A)mTq2 and KIF3B(S640A)mTq2 constructs. In addition, we also made KIF3B(S631D)mTq2, KIF3B(S640D)mTq2 and KIF3B(S631D and S640D)mTq2 double mutant constructs **(Figure S5)**. We then studied the interactions between these constructs and 14-3-3εmCit in the absence and presence of insulin, by estimating the proportion of blue and green pixels. We found no significant change between the wild type and S631A as well as S640A variants in absence of insulin (Figure 5A). In the presence of insulin, the interaction between KIF3B(S631A)mTq2 and 14-3-3εmCit increased significantly, whereas it remained unaltered between KIF3B(S640A)mTq2 and 14-3-3εmCit **(Figure 5B, Figure S6A,B)**. Together, the data suggested that S640 residue could play a significant role in interaction with 14-3-3ε. To test whether potential phosphorylation of the residue due to the insulin stimulation could enhance the interaction, we used the aspartate variants of these two residues. Although individual S631D and S640D variants did not show any significant change in absence of insulin, we found a significant increase in the proportion of blue and green pixels corresponding to an interaction, in the presence of S631D and S640D double aspartate (SS>>DD) mutant **(Figure 5C, Figure S7A)**. A similar result was obtained in the absence of insulin using AP-FRET, where KIF3B(SS>DD)mTq2 showed a significant increase in the FRET efficiency as compared to wild type or single aspartate mutants **(Figure S8)**. Furthermore, the lifetime values of none of the aspartate mutants changed in the presence of insulin **(Figure 5D and Figure S8B-G)**. Altogether, these results indicated that a potential phosphorylation of either S640 and/or S631 after insulin stimulation is likely to enhance the 14-3-3ε association with the KIF3B tail. These KIF3B mutant data further supported the previous conclusion of a direct association between 14-3-3ε and KIF3B tail *in vitro*.

**Figure 5.**
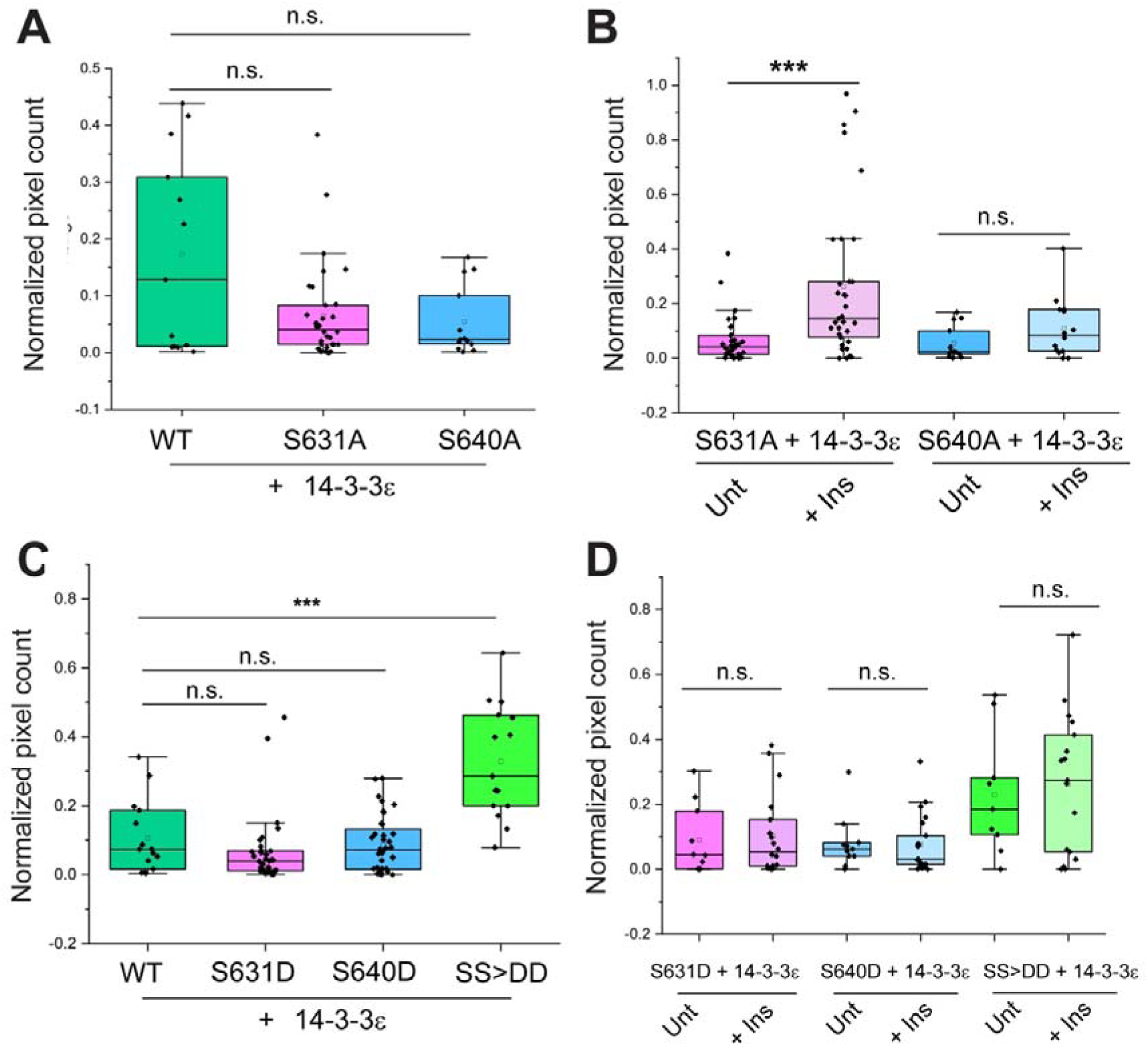
Effects of mutations in the serine residues, S631 and S640, respectively, within the predicted 14⍰3⍰3ε binding sites of the KIF3B tail. **A-D.** Box and whiskers plots show the fraction of pixels with the fluorescence lifetime of KIF3BmTq2 in the range of 3000–3800 ps (blue-green band) for each of the indicated conditions along the x-axis. The statistical significance of difference was estimated using Mann-Whitney U test. * Indicates p-value < 0.05 and *** indicates p-value < 0.001. Note that the alanine substitutions in the S631 and S640 residues indicated as S631A and S640A, respectively, does not alter the proportion of blue-green pixels significantly without the insulin stimulation (**A**). However, in the presence of insulin the proportion increases significantly in the S631A background, whereas it remained unchanged in the S640A background, indicating that S640 could play a significant role in binding 14-3-3ε (**B**). Single aspartate substitutions of the same residues, S631D and S640D, also did not show any increase in blue-green pixels with respect to the wild type KIF3BmTq2. However, the double substitution SS>DD, showed a significant increase in the blue-green pixels, indicating that phosphorylation of these two serine residues could play a significant role in 14-3-3ε interaction (**C**). The proportion of blue-green pixels did not increase any further in all the three backgrounds after insulin stimulation, suggesting that insulin stimulation could induce phosphorylation of one or both the serine residues (**D**).

### Kinesin-2 and 14-3-3ε interacting compartment are marked by Rab7

Finally, to understand the nature of the interacting compartments, we performed immunostaining on the cells used for split-APEX2 assay. Since we observed perinuclear localization of the resorufin signal from 14-3-3ε-S1 and KIF3B-S2, we asked whether this interaction might be associated with some of the Golgi compartments. To test this, we performed the AUR assay in cells coexpressing 14-3-3ε-S1 and KIF3B-S2 and co-stained them with a known Golgi marker, GM130. The nature of the signal obtained from anti-GM130 staining and AUR assay appeared very different, and we did not see any obvious colocalization between them **(Figure S9A)**. Next, we checked whether the signal associate with Rab4 vesicles, given the role of kinesin-2 in Rab-4 associated Glut4 vesicles upon insulin signaling^6,16,17^. While there were a few resorufin blobs colocalizing with Rab4 immunostaining, a majority of the Rab4 puncta did not colocalize with the resorufin signal **(Figure S9B)**.

To check whether 14-3-3ε is involved in kinesin-2 mediated transport of late endosomal vesicles^44^, we performed Rab7 antibody staining on the cells used for the AUR assay. We observed that almost all the blobs of the resorufin signal were marked by the Rab7 staining **(Figure S9C)**. This indicates a possible involvement of Kinesin-2 and 14-3-3ε on Rab-7 associated late endosomal vesicles, which are transported to cell periphery by kinesin-2^44^.

## Discussion

Heterotrimeric kinesin-2 transports cargoes to different cellular compartments, including cilia, axons and dendrites of neurons, and participates in retrograde transport from the Golgi apparatus to the endoplasmic reticulum^45^. Although heterotrimeric kinesin-2 is classically associated with intraflagellar transport ^46–48^, its involvement in other intracellular trafficking pathways suggests that its association with specific cargoes and regulatory proteins may be context-dependent. Previous studies have demonstrated that 14-3-3 proteins regulate microtubule-based transport through their interactions with kinesin motors and motor-associated proteins, including kinesin light chain 2, KIF1C, GRIP1-associated transport machinery, and the NudE/14-3-3 regulatory pathway^11,12,14,15,49^. These studies provide a precedent for the regulation of kinesin-dependent transport by 14-3-3 proteins. In the present study, we identified a previously uncharacterized, insulin-responsive association between the kinesin-2 motor subunit KIF3B and the regulatory protein 14-3-3ε in HEK293T cells. The spatially restricted and highly dynamic nature of this association argues against a constitutive interaction and is consistent with a context-dependent regulatory mechanism.

Using split-APEX2 and FLIM-FRET assays, we demonstrated that KIF3B and 14-3-3ε associate in spatially restricted subcellular regions, predominantly in the perinuclear region. Insulin stimulation increased the proportion of pixels exhibiting reduced donor fluorescence lifetimes, corresponding to increased FRET between KIF3B-mTq2 and 14-3-3ε-mCit. This increase was particularly evident in the population of pixels exhibiting weak to moderate FRET. Since the FRET signal was restricted to a subset of intracellular regions, averaging fluorescence lifetimes across the entire cytoplasm may underestimate the extent of the interaction within these regions. Time-resolved FLIM-FRET measurements further revealed the dynamic nature of this association, with the interaction-associated compartments appearing and disappearing over time. Importantly, insulin stimulation did not produce a comparable increase in FRET between KIF3B-mTq2 and the binding-deficient 14-3-3ε(K50E)mCit mutant. Similarly, mock treatment did not produce appreciable changes in the fluorescence lifetime distribution. These observations suggest that the insulin-dependent increase in FRET is not simply a consequence of changes associated with the imaging procedure or a generalized increase in molecular proximity.

The specificity of the observed association is further supported by the mutational analyses of both KIF3B and 14-3-3ε. Mutation of the conserved K50 residue in the ligand-binding groove of 14-3-3ε diminished the interaction signal, while mutations of the predicted interacting residues in the KIF3B tail altered its responsiveness to insulin stimulation. These observations are difficult to explain solely by nonspecific molecular crowding or co-compartmentalization arising from overexpression.

Nevertheless, since both split-APEX2 and FRET are proximity-based methods, they cannot independently establish direct physical binding between KIF3B and 14-3-3ε. Thus, while the combined experimental evidence supports a specific, insulin-responsive association between these proteins in cells, the possibility that additional cellular factors participate in stabilizing or mediating this association cannot be excluded.

Post-translational modifications, particularly phosphorylation, are common mechanisms regulating motor activity and association with cargo molecules (reviewed in ^5^). Several signaling pathways regulate kinesin-2-mediated transport through phosphorylation-dependent mechanisms ^50–53^. In the present study, we investigated two predicted 14-3-3-binding residues, S631 and S640, located in the C-terminal tail domain of KIF3B. Both FLIM-FRET and AP-FRET analyses revealed an increase in the association between KIF3B and 14-3-3ε with the double phosphomimetic mutant KIF3B(S631D, S640D), whereas the individual aspartate substitutions did not significantly enhance the basal interaction. Interestingly, the double phosphomimetic mutant exhibited elevated FRET signals even in the absence of insulin stimulation, and further insulin treatment did not produce an appreciable increase in the interaction signal. This suggests that the double phosphomimetic substitutions may partially mimic the insulin-stimulated state of KIF3B.

In contrast, neither of the individual alanine substitutions, S631A or S640A, produced a statistically significant reduction in the basal proportion of pixels exhibiting weak to moderate FRET compared with wild-type KIF3B, although a decreasing trend was observed. However, the two mutants responded differently to insulin stimulation. While KIF3B(S631A) retained the insulin-induced enhancement of the association, this response was largely abolished in KIF3B(S640A). These observations suggest that S640 plays an important role in regulating the insulin-responsive association between KIF3B and 14-3-3ε, whereas S631 may contribute to the interaction through a cooperative mechanism. The stronger effect observed with the double phosphomimetic mutant is consistent with the possibility that multiple residues act cooperatively rather than independently. Such a mechanism has been reported for several other 14-3-3-binding proteins, in which multiple phosphorylation sites contribute to recognition and binding by 14-3-3 proteins ^54–56^.

However, phosphomimetic substitutions do not necessarily reproduce the structural and biochemical consequences of phosphorylation. Therefore, the mutational analyses presented here do not directly demonstrate phosphorylation of KIF3B at S631 or S640. Although the findings are consistent with a phosphorylation-dependent regulatory mechanism, direct biochemical identification of phosphorylation at these sites, using approaches such as mass spectrometry or phospho-specific antibodies, would be necessary to establish this mechanism conclusively.

Both S631 and S640 are evolutionarily conserved from *Chlamydomonas* to humans, suggesting potential functional importance. Interestingly, a serine residue in the *Chlamydomonas* kinesin-2 orthologue CrFLA8, homologous to S631 of mammalian KIF3B, has been implicated in CDPK1/CaMKII-dependent phosphorylation associated with ciliary growth and regulation ^53,57^. This finding provides a precedent for phosphorylation-dependent regulation of the kinesin-2 motor. However, whether the conserved residues of mammalian KIF3B are phosphorylated in response to insulin stimulation remains to be established. The highly dynamic nature of the KIF3B–14-3-3ε association also raises the possibility that a balance between kinase and phosphatase activities regulates its formation and dissociation. Furthermore, 14-3-3 proteins have been reported to protect phosphorylated residues on their interacting partners from dephosphorylation by phosphatases^58^. Whether a similar mechanism contributes to the regulation of KIF3B remains an interesting possibility for future investigation.

We further investigated the subcellular localization of the KIF3B–14-3-3ε association using the split-APEX2 assay in combination with immunostaining for intracellular compartment markers. A significant proportion of the split-APEX2-positive puncta overlapped with Rab7-positive structures, suggesting that the association occurs, at least in part, within Rab7-positive endosomal compartments. Since Rab7 is commonly associated with late endosomal trafficking ^59–61^, this observation raises the possibility that the association participates in a regulated trafficking-related process. However, the overlap with Rab7 does not establish the precise identity of these compartments or demonstrate that the KIF3B–14-3-3ε association regulates their transport. More detailed characterization of the compartment and its associated cargoes will be necessary to understand the biological significance of this spatially restricted association.

A previous study from our laboratory demonstrated that heterotrimeric kinesin-2 participates in the transport of Rab4-positive vesicles towards axon terminals ^62^. Additionally, insulin stimulation in cultured adipocytes activates Rab4 GTPase, increases its association with kinesin-2, and enhances kinesin-2-dependent trafficking of GLUT4 vesicles towards the cell periphery ^6^. Insulin stimulation also promotes GLUT4 trafficking through the recruitment of 14-3-3 proteins to the RabGAP AS160 ^17^. Together, these findings suggest a possible functional connection between kinesin-2, 14-3-3 proteins, and insulin-regulated intracellular trafficking.

One hypothesis arising from these observations is that insulin signaling may facilitate the formation of a regulatory complex involving KIF3B, 14-3-3ε, AS160, and Rab4, thereby contributing to GLUT4 vesicle trafficking. However, this model remains speculative, as the present study does not demonstrate the formation of such a complex or investigate GLUT4 transport. Furthermore, the association observed in our experiments was localized, at least in part, to Rab7-positive structures rather than being directly linked to Rab4-positive vesicles. Therefore, the relationship between the Rab7-associated KIF3B–14-3-3ε interaction identified here and the previously reported Rab4-dependent trafficking pathways remains unclear. Future experiments examining the effect of selectively disrupting the KIF3B–14-3-3ε association on cargo localization and intracellular trafficking would help establish whether this interaction has a functional role in insulin-responsive transport.

Taken together, our findings identify a previously uncharacterized, spatially restricted, and insulin-responsive association between KIF3B and 14-3-3ε in mammalian cells. The combined evidence from split-APEX2, AP-FRET, live-cell FLIM-FRET, and mutational analyses supports the specificity of this association and implicates the C-terminal tail domain of KIF3B, particularly the region containing S640, in its regulation. Although the molecular mechanism and downstream physiological consequences remain unresolved, identifying a stimulus-responsive association between a kinesin motor subunit and a regulatory protein provides a foundation for investigating additional mechanisms controlling kinesin-2-mediated intracellular transport.

### Limitations and Future Scope of the Study

The present study has several limitations that should be considered when interpreting the findings.

First, although the association between KIF3B and 14-3-3ε was supported by multiple cell-based assays, the quantitative imaging experiments were performed using transiently expressed, fluorescently tagged proteins. Such expression systems can alter the relative abundance, localization, and proximity of proteins and may not fully reflect their behaviour at endogenous levels. Although, the use of binding-deficient and phosphosite mutants provides evidence against a purely nonspecific overexpression-associated signal, the possibility of an indirect association mediated by additional cellular proteins remains unresolved. Future studies using endogenous protein tagging, proximity ligation assays, or optimized biochemical approaches could help address these limitations.

Second, the assays employed in this study have inherent methodological constraints. Split-APEX2 was selected because enzymatic signal amplification permits detection of weak or transient protein associations. However, its requirement for H₂O₂ treatment introduces the possibility of oxidative perturbations, and complementation-based proximity assays may be influenced by local protein concentrations. Similarly, AP-FRET measurements performed in fixed cells may underestimate transient interactions if fixation alters protein organization. Live-cell AP-FRET also presents challenges because diffusion of unbleached acceptor molecules into the photobleached region can affect FRET estimation. Therefore, greater emphasis was placed on live-cell FLIM-FRET measurements, which do not require acceptor photobleaching and can monitor changes in fluorescence lifetime during insulin stimulation. Importantly, the insulin-dependent changes observed using FLIM-FRET occurred without H₂O₂ treatment, providing evidence independent of potential oxidative effects associated with split-APEX2. Nevertheless, FRET measurements report molecular proximity rather than direct binding, and their interpretation may be influenced by fluorophore orientation and the heterogeneity of interacting and non-interacting populations.

Third, although we found that the insulin-responsive KIF3B–14-3-3ε association occurred in spatially restricted regions overlapping with Rab7-positive structures, the identity and biological function of these compartments remain incompletely characterized. The present study did not establish whether the association influences motor activity, cargo recruitment, vesicle trafficking, or other cellular processes. An attempt to identify the potential interaction partners and associated cargoes using split-APEX2-based proximity labelling was unsuccessful because of the low yield of biotin-labelled proteins. Optimization of the assay using a different cell line, improved labelling conditions, or a larger culture system may help identify additional proteins associated with this interaction. Such experiments could also provide insight into the functional significance of the Rab7-positive compartments.

Fourth, there are seven isoforms of 14-3-3 proteins in mammals. The central amphipathic groove through which 14-3-3 dimers recognize their target proteins is highly conserved across isoforms ^54,56^. This conservation contributes to overlapping target recognition and some functional redundancy among different isoforms.

However, the mechanisms underlying isoform-specific target recognition remain incompletely understood. Since the present study examined only 14-3-3ε, it remains possible that KIF3B also associates with other 14-3-3 isoforms. A comparative analysis of these isoforms could help determine whether the observed association is specific to 14-3-3ε or represents a broader mechanism of kinesin-2 regulation.

Finally, although our mutational analyses indicate that the region containing S631 and S640, particularly S640, contributes to the insulin-responsive association between KIF3B and 14-3-3ε, the phosphorylation status of these residues has not been directly determined. Furthermore, the upstream kinases and phosphatases potentially responsible for regulating the association remain unidentified. Future phosphoproteomic analyses, targeted biochemical experiments, and identification of proteins in the vicinity of the KIF3B–14-3-3ε complex may help establish the signaling mechanism responsible for its regulation.

Despite these limitations, the present study provides evidence for a specific, dynamically regulated association between KIF3B and 14-3-3ε in mammalian cells. Further investigation of its molecular mechanism, associated cargoes, and physiological consequences will be necessary to determine whether this association contributes to the regulation of kinesin-2-dependent transport in response to cellular signaling.

## Materials and Methods

### 1. Cloning of constructs in mammalian expression vectors

MmKIF3A and MmKIF3B clones, obtained Dr. Martin Engelke (University of Illinois), and human 14-3-3ε clone, obtained from Dr. Sorab Dalal (ACTREC, Kharghar, India), were used to amplify and subclone respective cDNAs into different vectors. For FRET constructs, KIF3A and 14-3-3ε were cloned in mCitrineN1 vector (Addgene, Plasmid#54594) between Xho1 and Kpn1. mTurquoise2 was subcloned from mKOkappa-2A-mTurquoise2 (Addgene #98837) into p220ME_MmKIF3B(fl)-mCherry using BamH1 and Not1 restriction sites which replaced the mCherry tag. Site directed mutagenesis, using overlapping PCR primers, were used to introduce K50E mutation in 14-3-3ε and other mutations (S631A, S640A, S631D, S640D and double mutants S631D with S640D, labelled as SS>>DD) in the C-terminal tail region of KIF3B clones. S1 and S2 fragments of ascorbate peroxidase 2 (APEX2) were amplified from mito-V5-APEX2 vector (Addgene, Plasmid#72480) and subcloned into p220ME_MmKIF3B(fl)-mCherry between BamH1 and Not1 for KIF3B-S2, and between Xho1 and Kpn1 of 14-3-3□-mCit for making 14-3-3□-S1. For the mutant 14-3-3□(K50E)-S1, single point mutation was inserted in 14-3-3□-S1 with site-directed mutagenesis. All clones were confirmed by sequencing for atleast two rounds.

### 2. Tissue culture

HEK293T cells were maintained in an incubator at 37°C with 5% CO_2_ in Dulbecco’s Modified Eagle Medium (DMEM, Gibco-Thermo Fisher Scientific) containing 10 % FBS (Gibco-Thermo Fisher Scientific) and 1X Penicillin-Streptavidin solution (Gibco-Thermo Fisher Scientific).

### 3. Split-APEX2 assay

2×10^5^ cells were seeded on a poly-D-lysine-coated coverslip and transfected with the S1 and S2 constructs after 24 hours, using Lipofectamine 3000 (Thermo Fisher Scientific), according to manufacturer’s protocol. Cells were allowed to grow for 48 hours post-transfection in at 37°C and 5% CO_2_. After 48 hours, the media was discarded by aspiration, and the cells were given two PBS washes. The cells were then treated with 50µM of Amplex Ultra Red (Thermo Fisher Scientific) with 0.02% H_2_O_2_ in PBS, and incubated in dark for 20 minutes at room temperature. Following which, the solution was discarded, cells were washed with PBS for two to three times, and incubated with ice-cold 4% paraformaldehyde (PFA, Sigma Chemical Co. USA) for 20 minutes at room temperature. After the PFA fixation, the cells were washed again with PBS for three times and mounted on a glass slide with a drop of Vectashield^TM^ (Vector Labs, Inc).

### 4. Fluorescence lifetime imaging (FLIM)-FRET

Approximately 2 × 10□ cells were seeded in 35-mm culture dishes containing complete DMEM media and maintained at 37°C in a humidified incubator with 5% CO₂. After 24 hours, cells were transfected with FRET expression constructs using Lipofectamine 3000, according to the manufacturer’s instructions and incubated under the same conditions. Forty to forty-eight hours post-transfection, the culture medium was removed, and cells were washed once with phosphate-buffered saline (PBS). Cells were subsequently immersed in 1.5 ml of imaging buffer (110 mM NaCl, 40 mM NaHCO_3_, 1 mM CaCl_2_, 5 mM KCl, 1mM MgCl_2_, 20 mM HEPES and 1 g/L D-glucose) and imaged using Zeiss LSM 980.

Fluorescence lifetime imaging microscopy was performed on an upright confocal microscope (Zeiss LSM 980) equipped with a Becker & Hickl fluorescence lifetime imaging module for time-correlated single photon counting (TCSPC). Confocal images were first acquired to assess the expression levels of mTq2 and mCit prior to lifetime measurements. Cells expressing both the fluorophores with mCit levels at least 1.5 times higher than that of mTq2 were selected for FLIM. Plan Apo 20x (N.A. 1.0) water dipping objective, was used to acquire all the confocal and corresponding lifetime images with the 512 x 512 pixels frame size, along with 3.5X digital zoom and at 16 bits per pixel.

Fluorescence lifetime measurements were performed using a Ti: Sapphire Mai Tai multiphoton laser operating at an excitation wavelength of 800 nm, with a repetition rate of 80 MHz. Fluorescence lifetime data were acquired for approximately 180 seconds per field of view and analysed using SPCImage® software. For analysis, the cytoplasmic region of each cell was manually selected as the region of interest (ROI). Auto-generated Instrument Response Function (IRF) from the SPCImage® software and a binning factor of 5 were used to fit the fluorescence decay curves to a single-exponential decay model. The resulting pixel-wise fluorescence lifetime values were used to generate frequency distribution histograms and determine the number of pixels within three lifetime ranges: 3000–3400 ps (blue), 3400–3800 ps (green), and 3800–4200 ps (red).

### 5. Acceptor photobleaching-F⍰rster Resonance Energy Transfer (AP-FRET)

For preparing sample for Acceptor photobleaching FRET (AP-FRET), 2 x 10^5^ cells were seeded on a poly-D-lysine-coated coverslip in a 35 mm dish. After 24 hours, cells were transfected with mTurquoise2 and mCitrine constructs using Lipofectamine 3000, as per the manufacturer’s protocol. Cells were then grown for 48 hours in the incubator at 37°C with 5% CO_2_. After 48 hours, the cells were taken out from the incubator, the media was discarded, and cells were washed with PBS. Cells were then fixed with ice-cold 4% PFA for 20 minutes at room temperature. Post-fixation, cells were washed thrice with PBS and the coverslip was then then mounted in vectashield on a glass slide.

AP-FRET experiments were performed using an Olympus FV3000 confocal microscope equipped with 445 nm and 514 nm solid-state lasers to excite mTurquoise2 and mCitrine, respectively. Fluorescence images were acquired using a 60X oil-immersion objective (NA 1.42) and a high-sensitivity spectral detector (HSD) with a gallium arsenide phosphide (GaAsP) photodetector. Emission windows of 460–500 nm and 527–567 nm were used to collect mTurquoise2 and mCitrine fluorescence, respectively. Pre-bleach images were acquired in both donor and acceptor channels, following which mCitrine was photobleached using a high-intensity 514 nm laser. During photobleaching, the detector voltage was set to its minimum to prevent detector saturation and potential damage. Subsequently, post-bleach images were acquired in both channels using the same imaging parameters as those used for the pre-bleach images.

For analysis, 3-5 random ROIs were drawn in the cytoplasm of the cell and both donor (mTq2) and acceptor (mCitrine) fluorescence intensities before and after acceptor photobleaching were quantified. FRET fraction was calculated using the following equation:

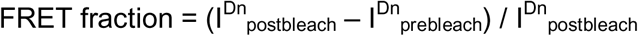

The FRET fraction was normalised to the bleaching fraction which was obtained using the formula:

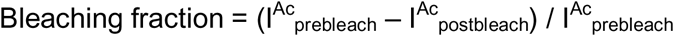

Hence, final FRET efficiency was calculated by:

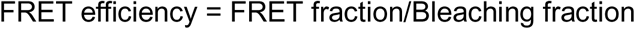

### 6. Insulin treatment of Cells

For the before-and-after insulin treatment experiments, ROIs containing one or more cells expressing the both the mTq2 and mCit tagged constructs were selected for imaging. The FLIM images were first acquired under untreated conditions. Subsequently, approximately 200 µL of imaging buffer was withdrawn from the dish without disturbing it, mixed with 15 μl of the 1.7 μM insulin stock solution, and added back to the dish to achieve a final insulin concentration of 17 nM. Fluorescence lifetime data were then acquired again immediately following the treatment from the same field of view.

For the time-course experiments, cells growing in DMEM complete media were pre-treated with 17 nM insulin and kept in the incubator at 37°C and 5% CO₂ for 5 minutes. The cells were subsequently washed with PBS containing 17 nM insulin and imaged in imaging medium supplemented with the same concentration of insulin for one hour at an interval of 5 minutes. Approximately 5-10 minutes were spent in finding appropriate frames for imaging, due to which the time-course data were always recorded 25 minutes after the insulin treatment.

For serum starvation experiments, cells were cultured in serum-free DMEM for 16 h prior to FLIM. Insulin treatment was given in the same fashion as described above.

### 7. Immunostaining

For immunofluorescence staining, cells were incubated in blocking and permeabilization buffer (B-PBS-T; phosphate-buffered saline containing 1% (w/v) bovine serum albumin (BSA) and 0.1% (v/v) Triton X-100) post-fixation for 1 hour at room temperature. Cells were then incubated with primary antibodies diluted in B-PBS-T buffer overnight at 4°C in a humidified chamber.

Following primary antibody incubation, the antibody solution was removed and cells were washed two to three times with B-PBS-T buffer. Cells were subsequently incubated with species-appropriate Alexa Fluor–conjugated secondary antibodies (anti-rabbit or anti-mouse) diluted in B-PBS-T buffer, along with Hoechst nuclear stain (10 μg/mL), for 1–2 hours at room temperature in the dark. After secondary antibody incubation, cells were washed once with B-PBS-T buffer followed by two washes with PBS. Coverslips were then mounted onto glass slides using Vectashield antifade mounting medium and stored 4°C until imaging.

### 8. Prediction of potential 14-3-3-binding sites

The online prediction tool, 14-3-3 Pred was used to identify potential 14-3-3-binding motifs on all the three subunits of kinesin-2. The proteins sequences corresponding to the following accession codes were used as input sequence: KIF3A (P28741); KIF3B (Q61771); and KAP3 (P70188), respectively. The sites having ANN, PSSM and SVM scores higher than 0.55, 0.8 and 0.25, respectively were identified as potential 14-3-3-binding sites.

### 9. Statistical analysis

Paired Student’s T-test was used to calculate the significance of difference for before and after Insulin treatment data. Welch’s unpaired t-test was used to calculate the significance of difference in the time-course experiment. Apart from that, non-parametric Mann-Whitney U test was performed to calculate significance of difference for all other datasets.

### 10. Accession codes

The UniProt primary accession ID of the proteins used in the study are: 14-3-3ε: P62258; MmKIF3A: P28741; MmKIF3B: Q61771

## Acknowledgement

We are grateful to Dr. Sorab Dala of ACTREC, Kharghar, India, for providing the 14-3-3ε constructs and extensive discussion during the project, and Dr. Martin Engelke, University of Illinois, for the plasmids p220ME_MmKIF3B(fl)-mCherry and p285ME_MmKIF3A(fl), which were used to sub-clone various plasmids containing MmKIF3B and MmKIF3A. We also acknowledge Dr. Raikamal Pal and Yukta Khurana, DBS, TIFR, Mumbai, for providing the split-APEX2 constructs, and other assistance during the project. Additionally, we acknowledge members of the Debasis Das’ lab and Prof. Ullas Kolthur’s labs at DBS, TIFR, Mumbai, for numerous assistances; Ms. Boby KV for help with the operations of Zeiss 980 fitted with the multiphoton laser and for training on the FLIM acquisition software. We would like to thank Prof. Sudipta Maiti and his students, Dr. Debsankar Roy and Dr. Arpan Dey, for their help in standardising the FLIM-FRET experiment and analysis. The project was funded through an intramural grant from TIFR, DAE, and an EMR grant from the SERB (No. CRG/2020/005396, dated 30-Nov-2021), Government of India to KR. DK and AT were supported by fellowships from TIFR, DAE, Government of India.

---

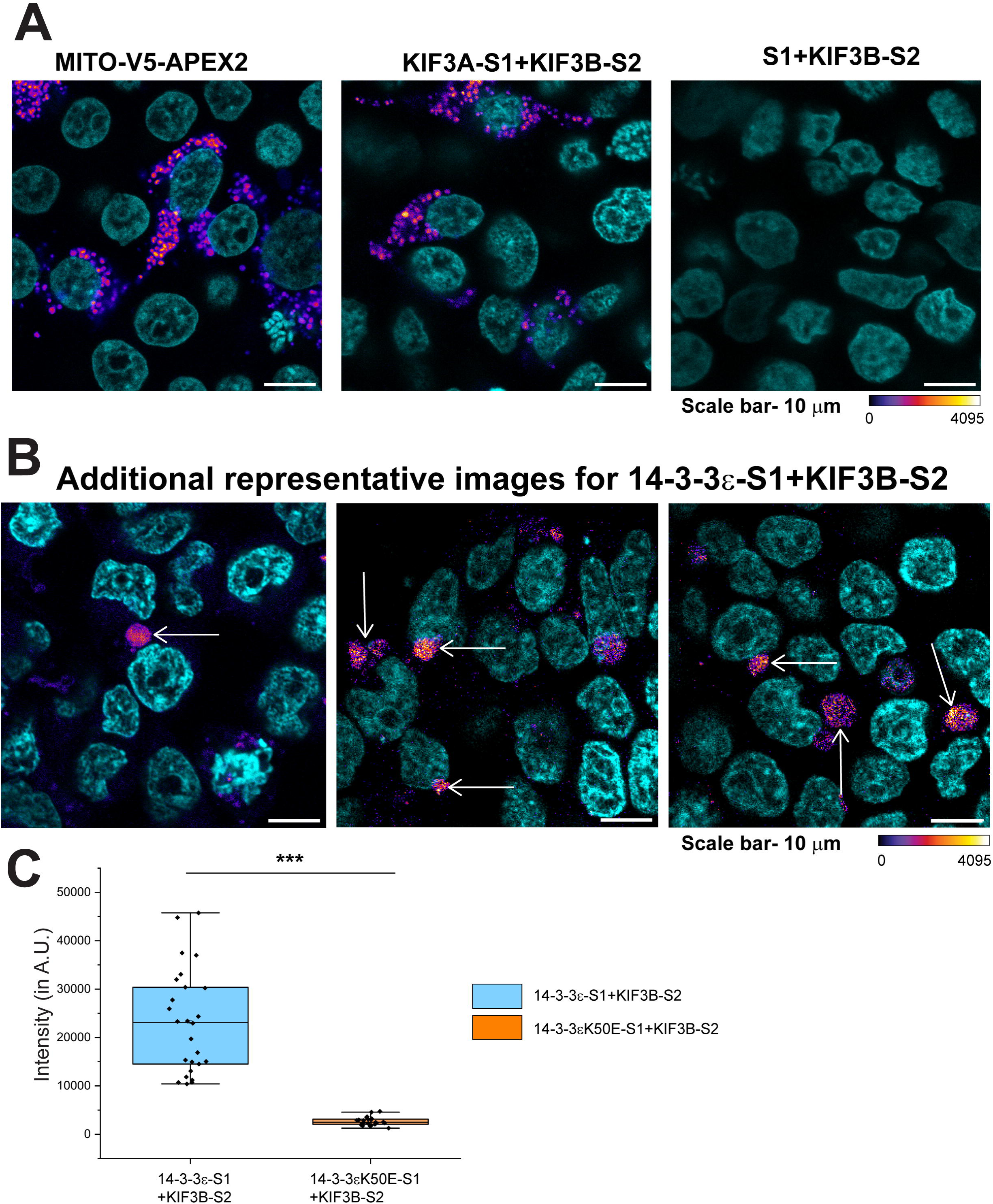

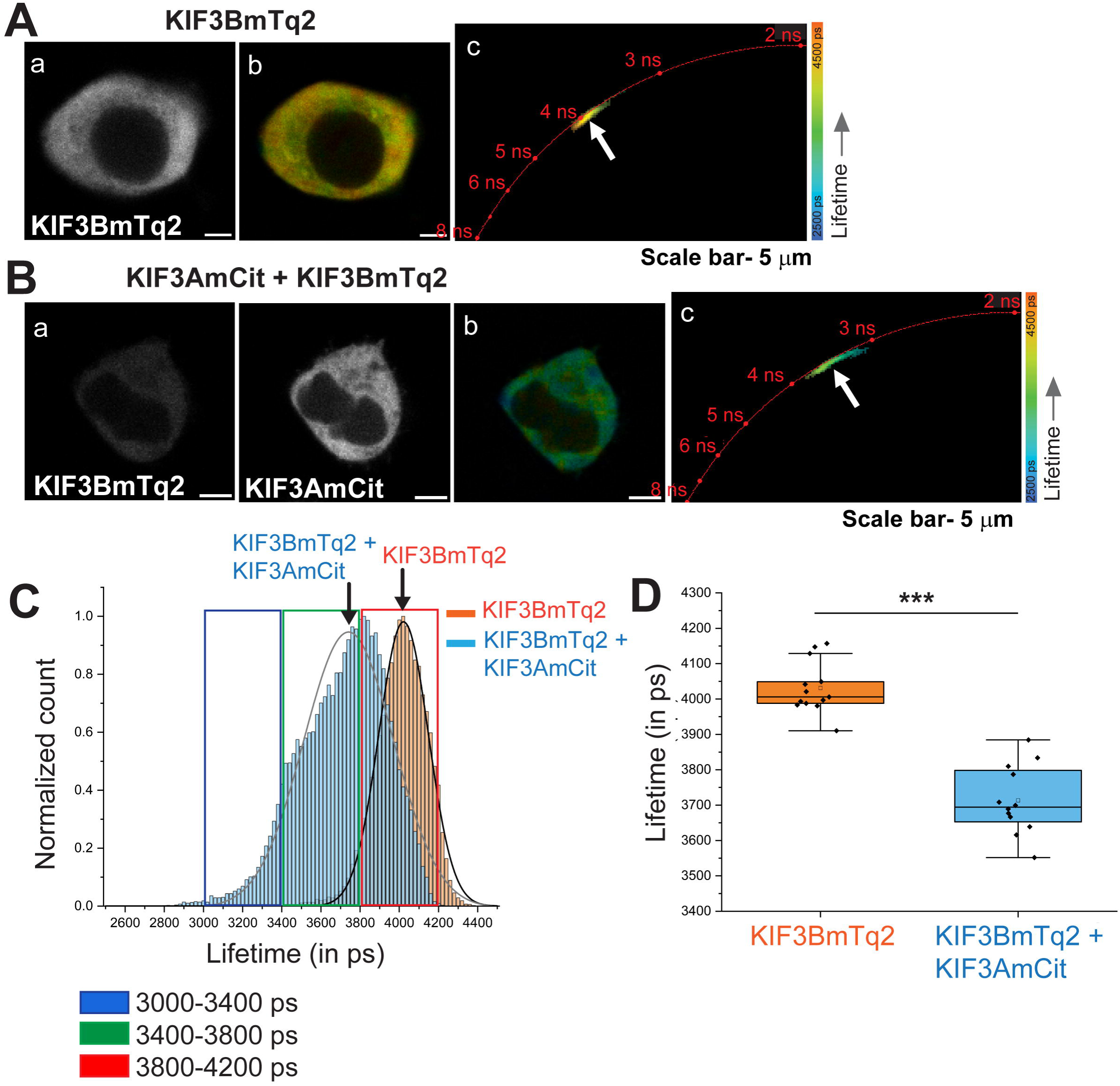

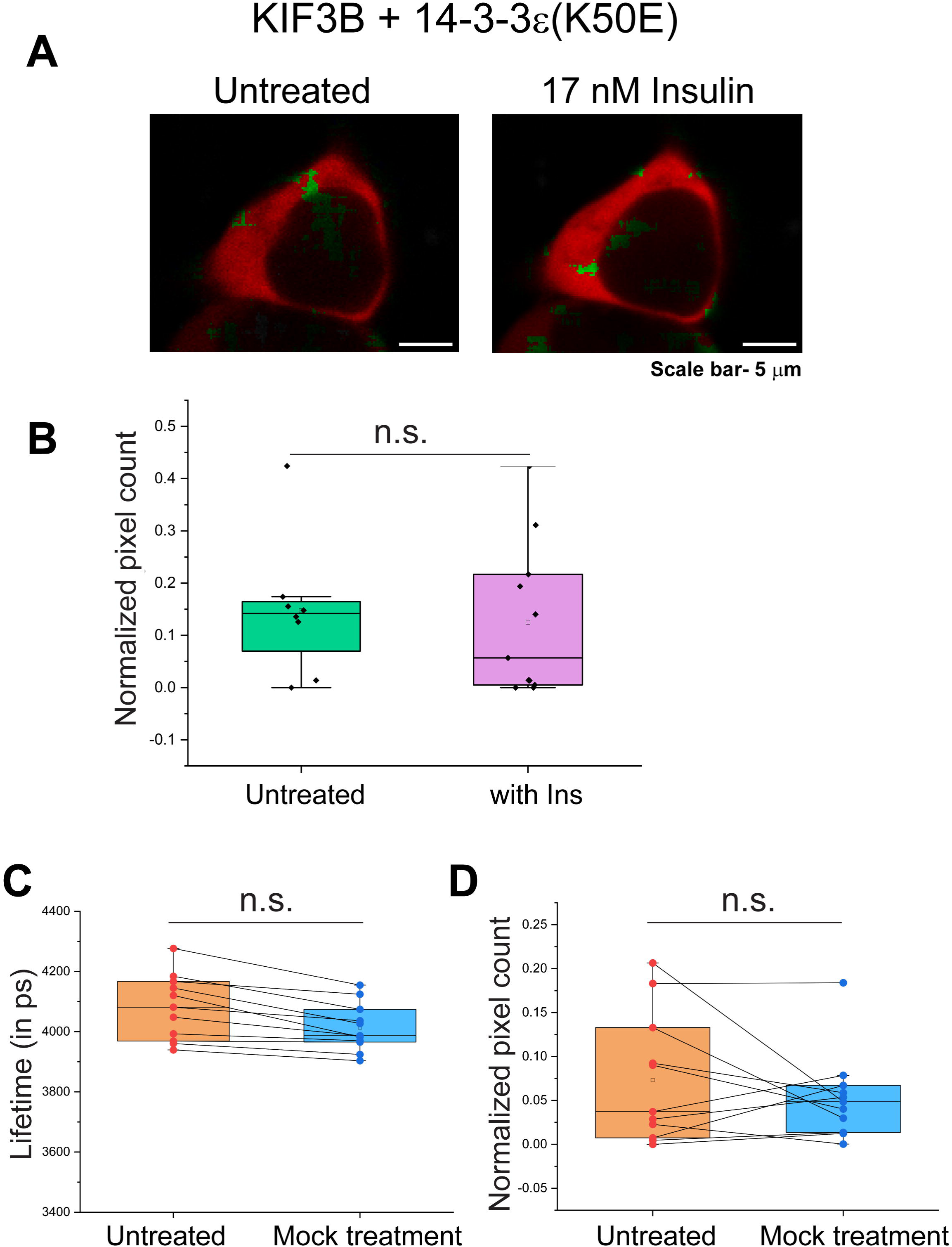

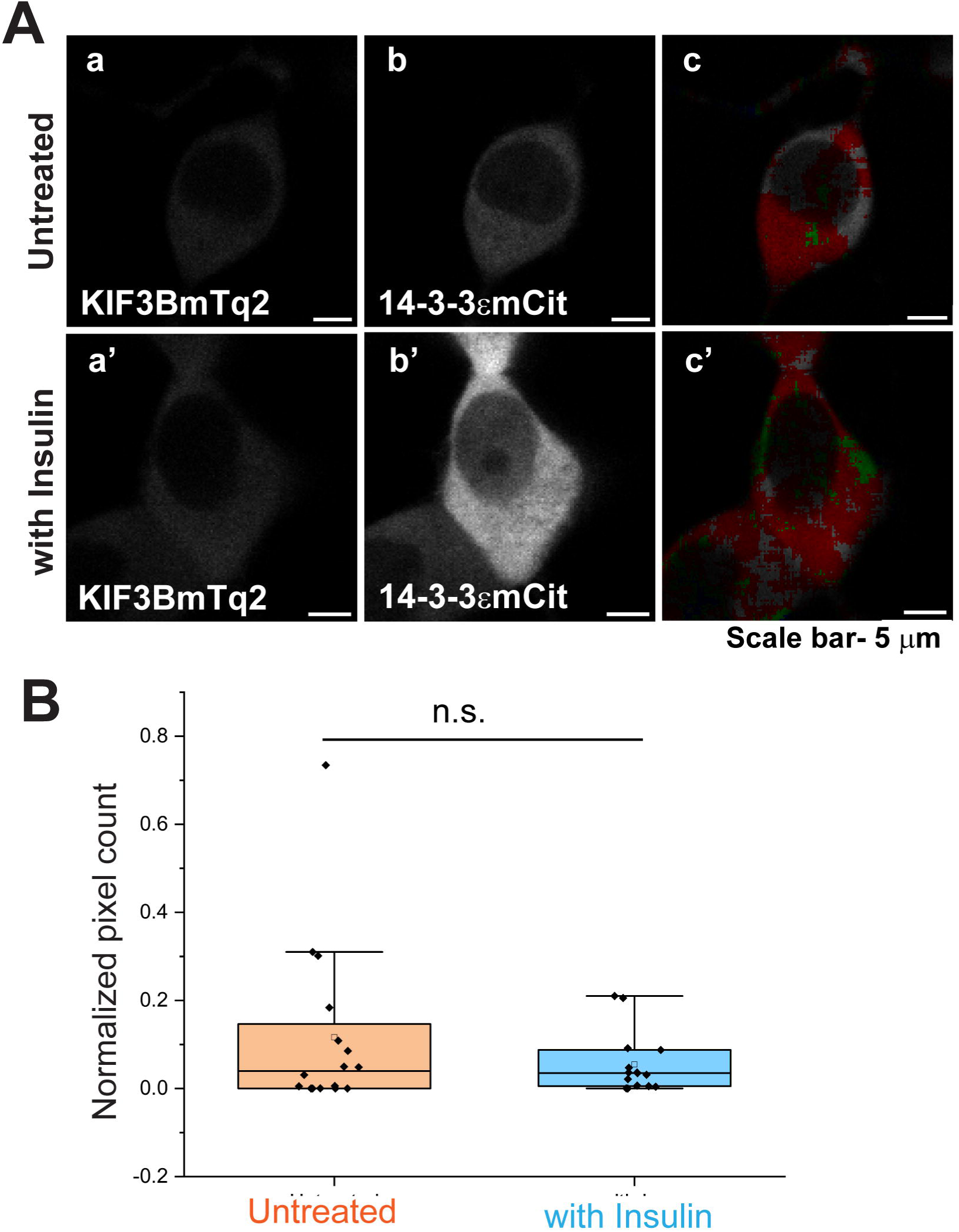

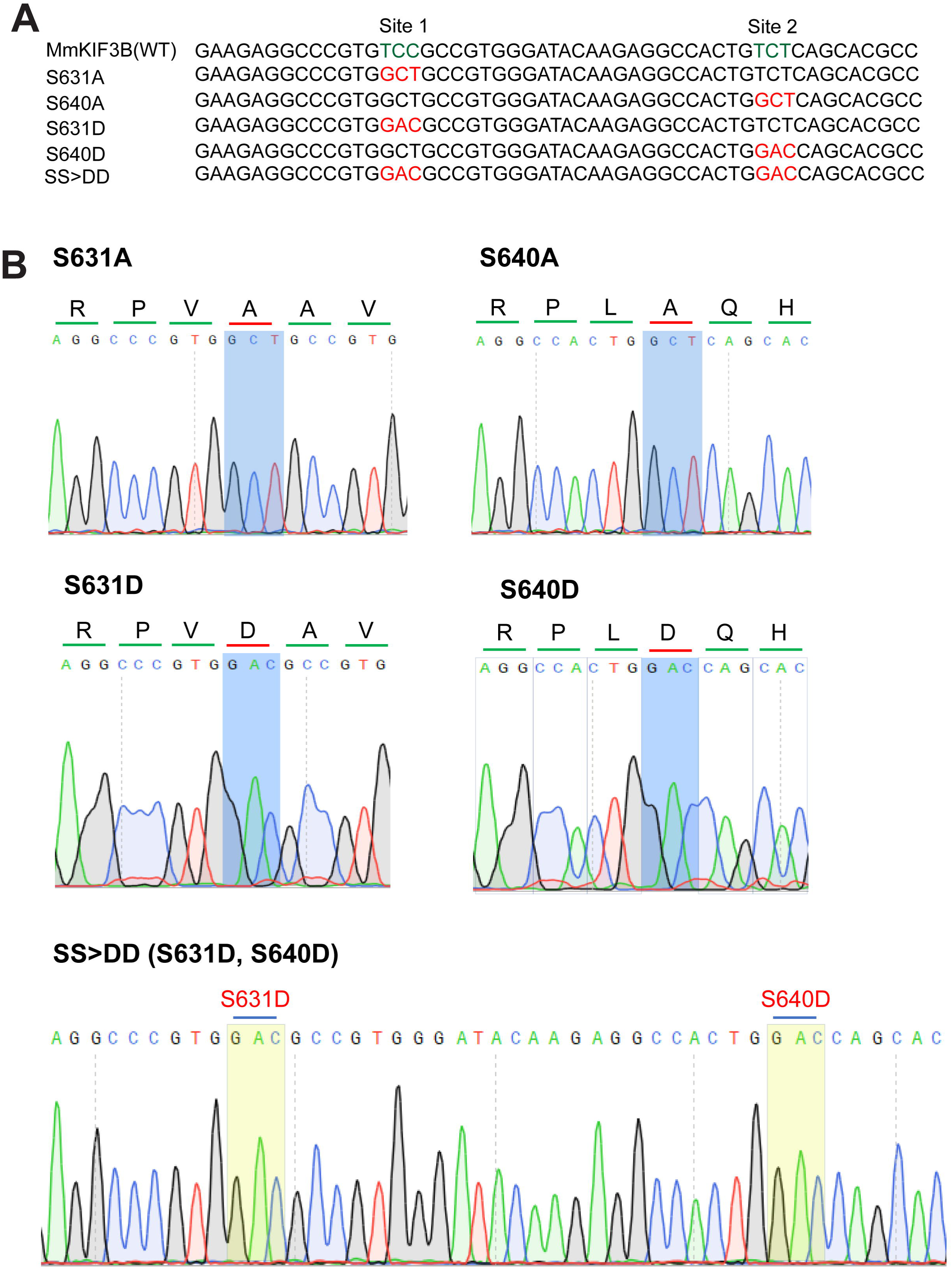

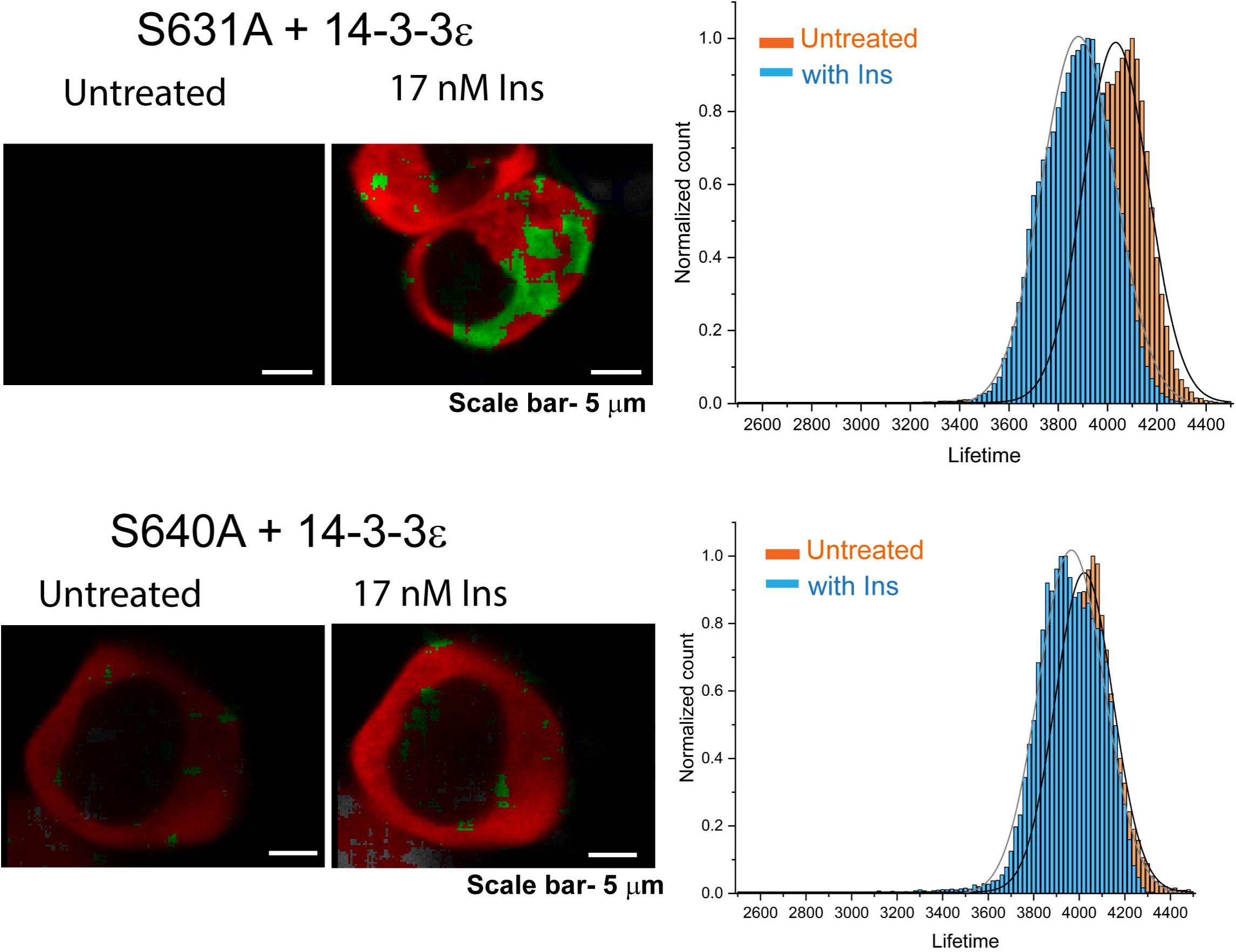

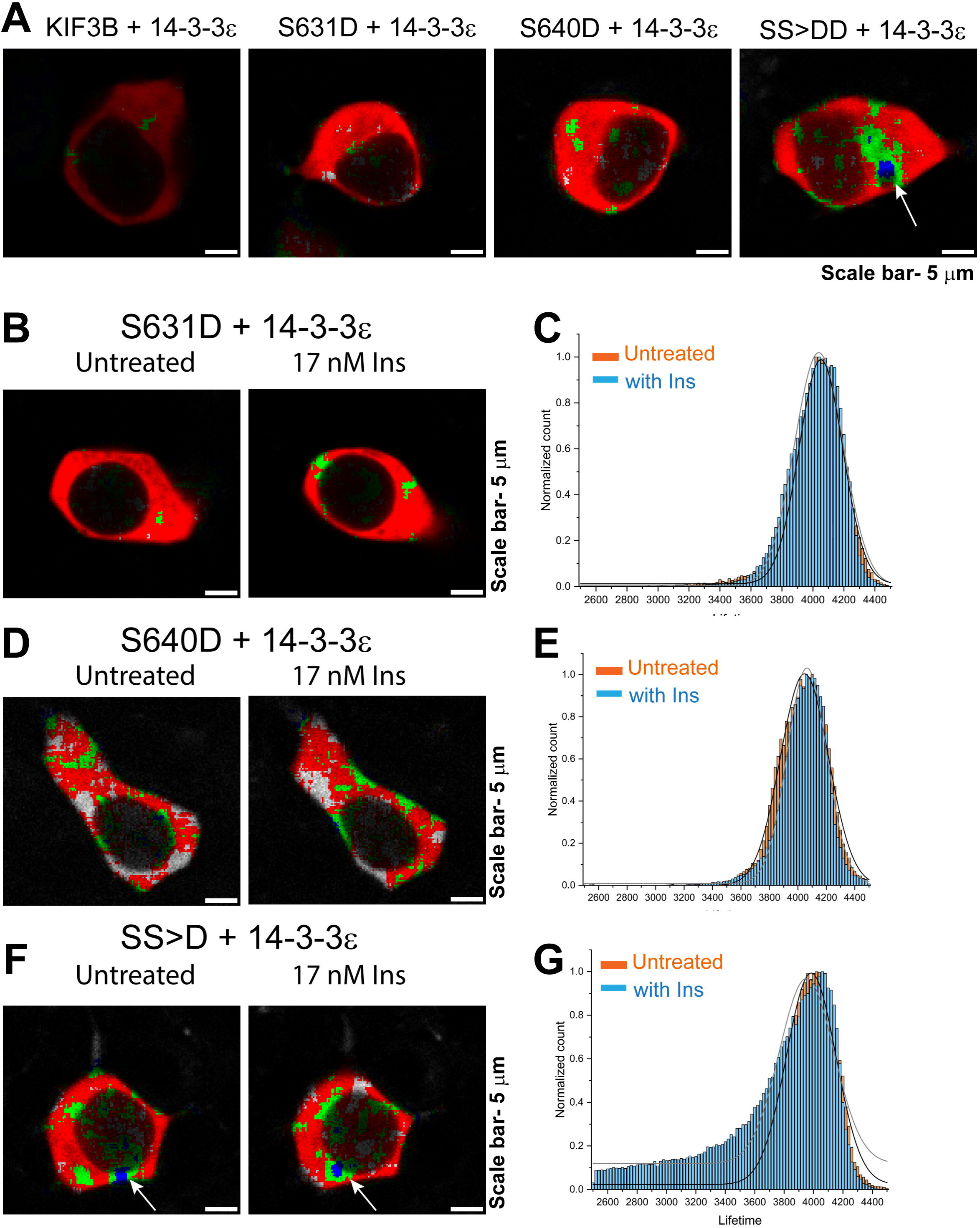

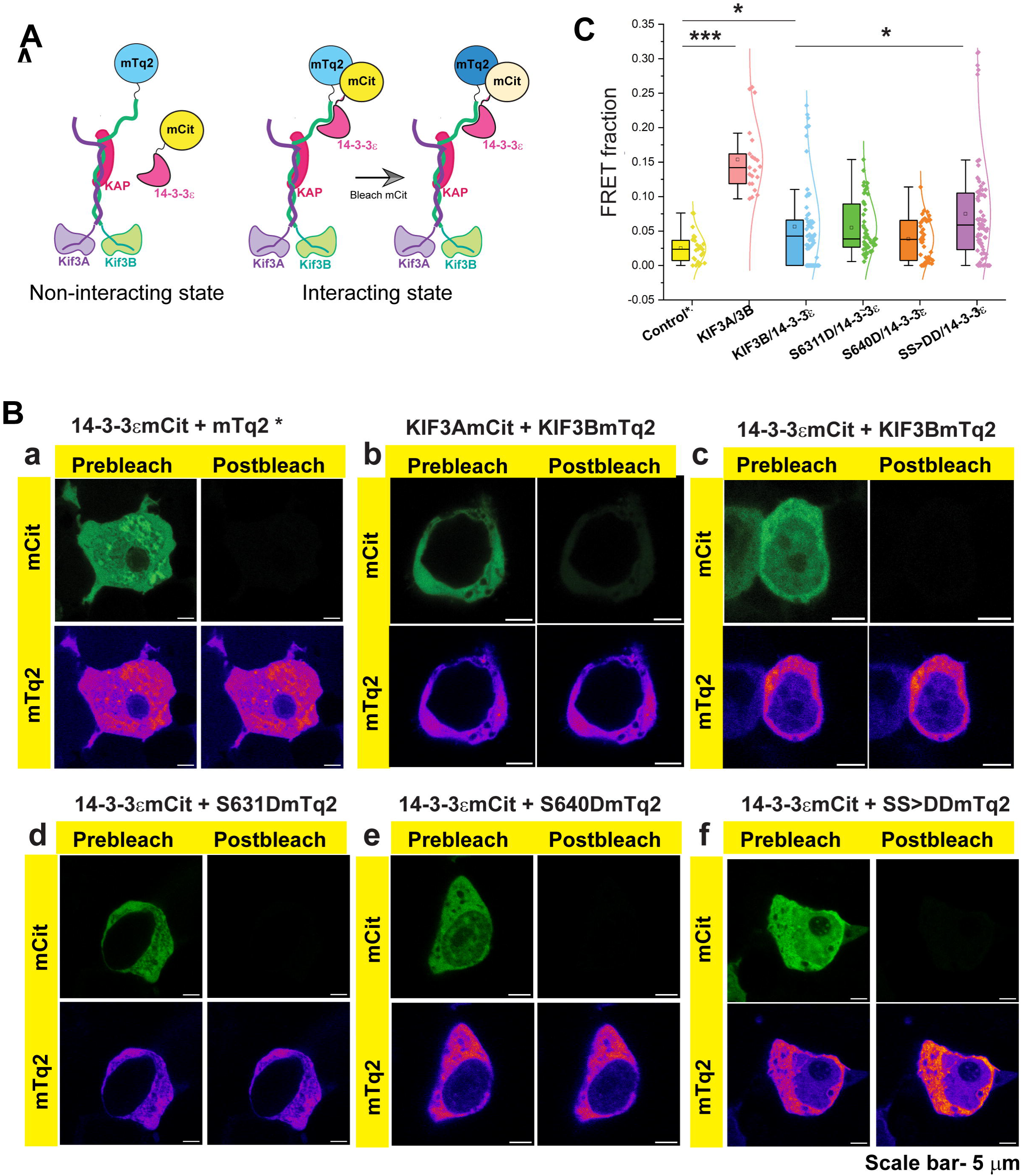

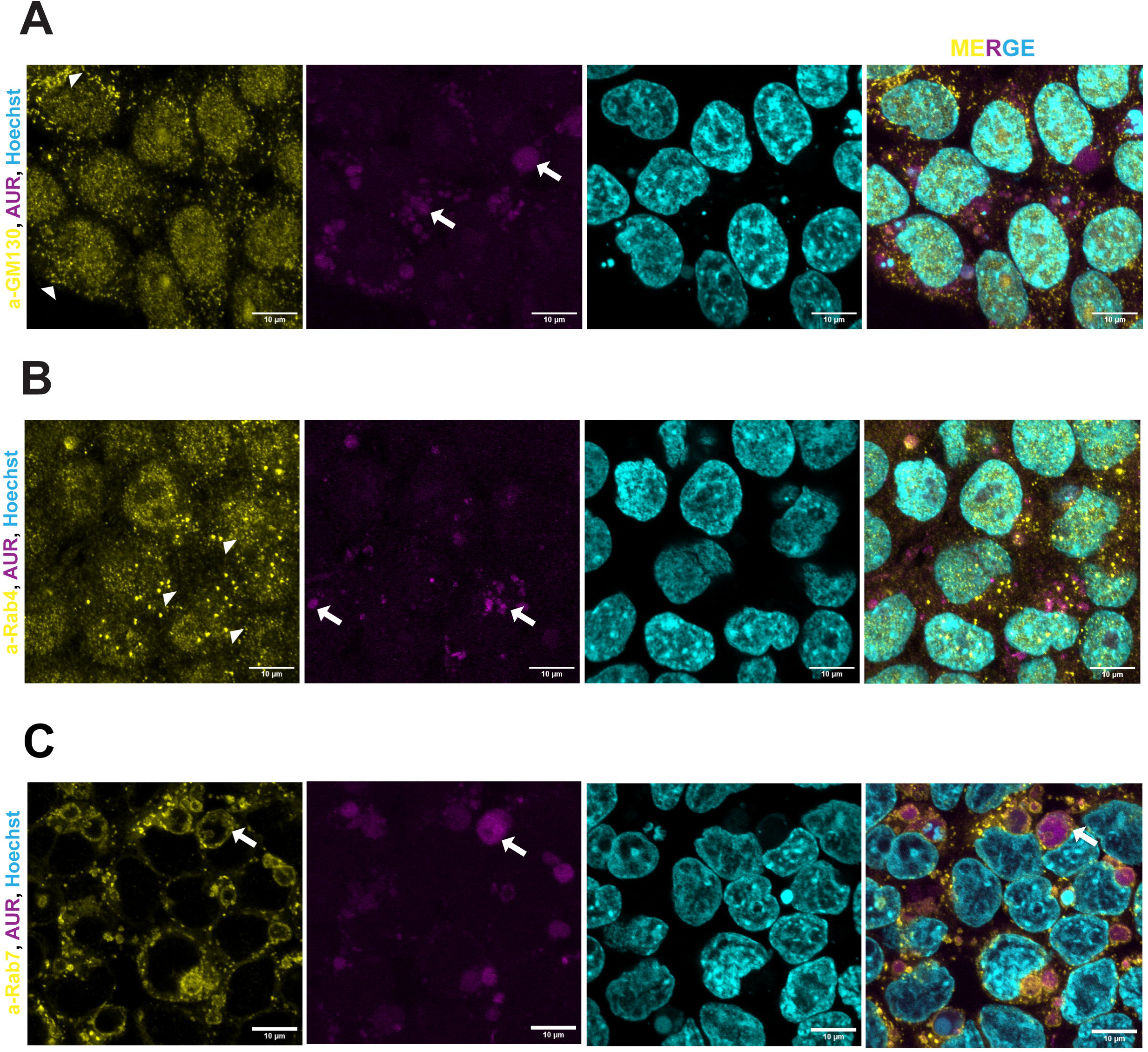

