## Supplementary information for "Insulin stimulation induces dynamic interaction between 14-3-3ε and the C-terminal tail domain of Kinesin-2 subunit KIF3B in vitro"

| **Serial no.** | **Supporting information** | **Number** |
| --- | --- | --- |
| 1. | Number of pages | 16 |
| 2. | Number of figures | 9 |
| 3. | Number of tables | 3 |

**Supplementary figure S1**

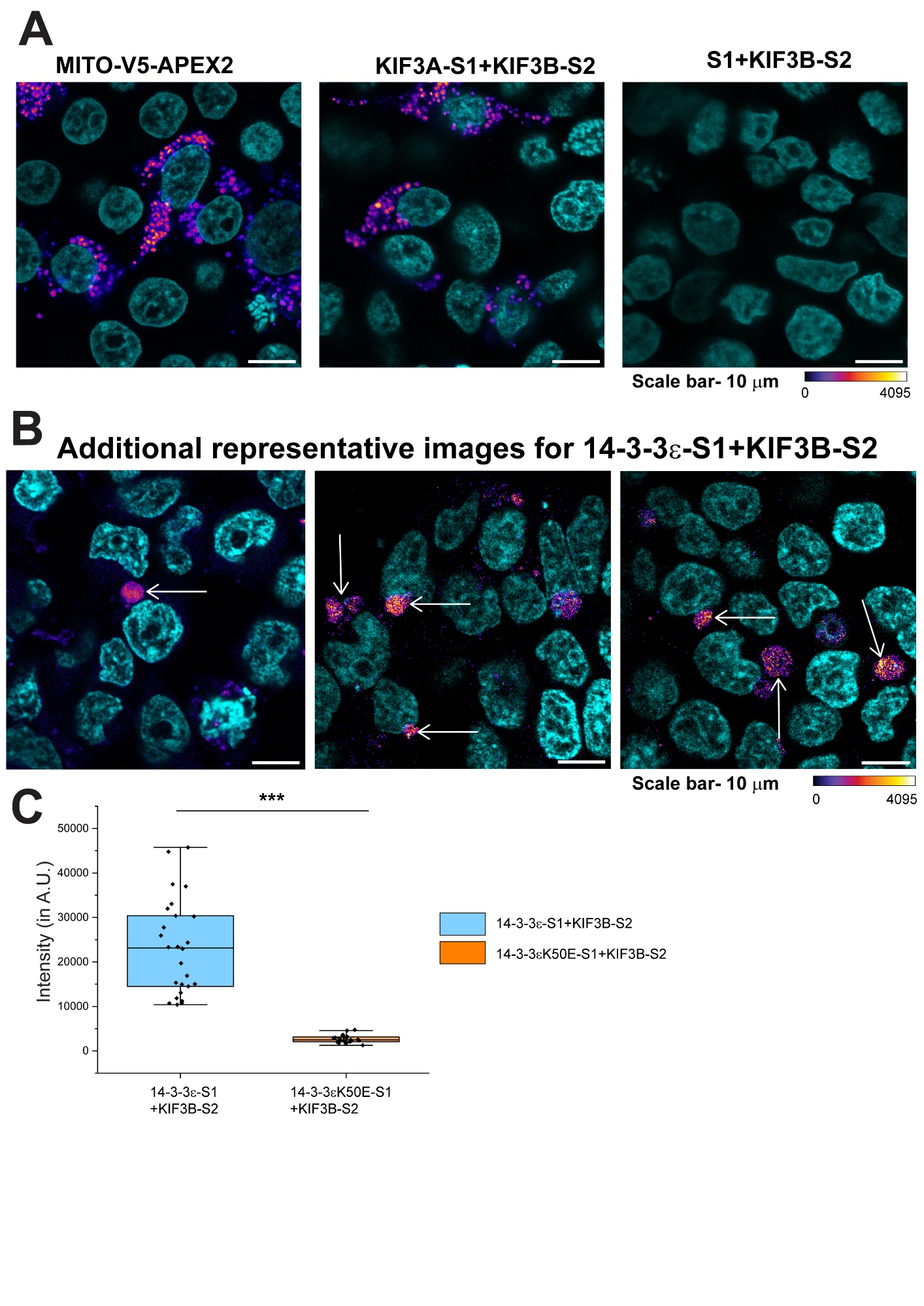

**Supplementary figure S1. Validation of split-APEX2 assay with full length mito-APEX2 and negative control pair.**

A. Split-APEX2 assay in cells transfected with Mito-V5-APEX2 (positive control); KIF3A-S1 and KIF3B-S2 (positive control); and untagged S1 fragment and KIF3B-S2, as indicated on top of each image. The resorufin signal is shown in false colour Fire LUT according to the calibration bar at the bottom of the panel. Nucei are stained with hoechst dye (in cyan). Scale bars ~ 10 μm
B. Additional representative images from 3 independent experiments, showing the interaction between 14-3-3ε-S1 and KIF3B-S2. Scale bars ~ 10 μm
C. Quantification of resorufin signal from co-transfection of 14-3-3ε-S1 and KIF3B-S2; and untagged S1 fragment and KIF3B-S2, as indicated on the x-axis. Each point represents an ROI drawn for 10 nuclei. The pair-wise significance of difference was estimated using Mann-Whitney U test, *** indicates p value < 0.001. N=2; n=25 ROIs (250 cells) for 14-3-3ε-S1 and KIF3B-S2 pair and n=21 ROIs (210 cells) for S1 and KIF3B-S2.

**Supplementary figure S2**

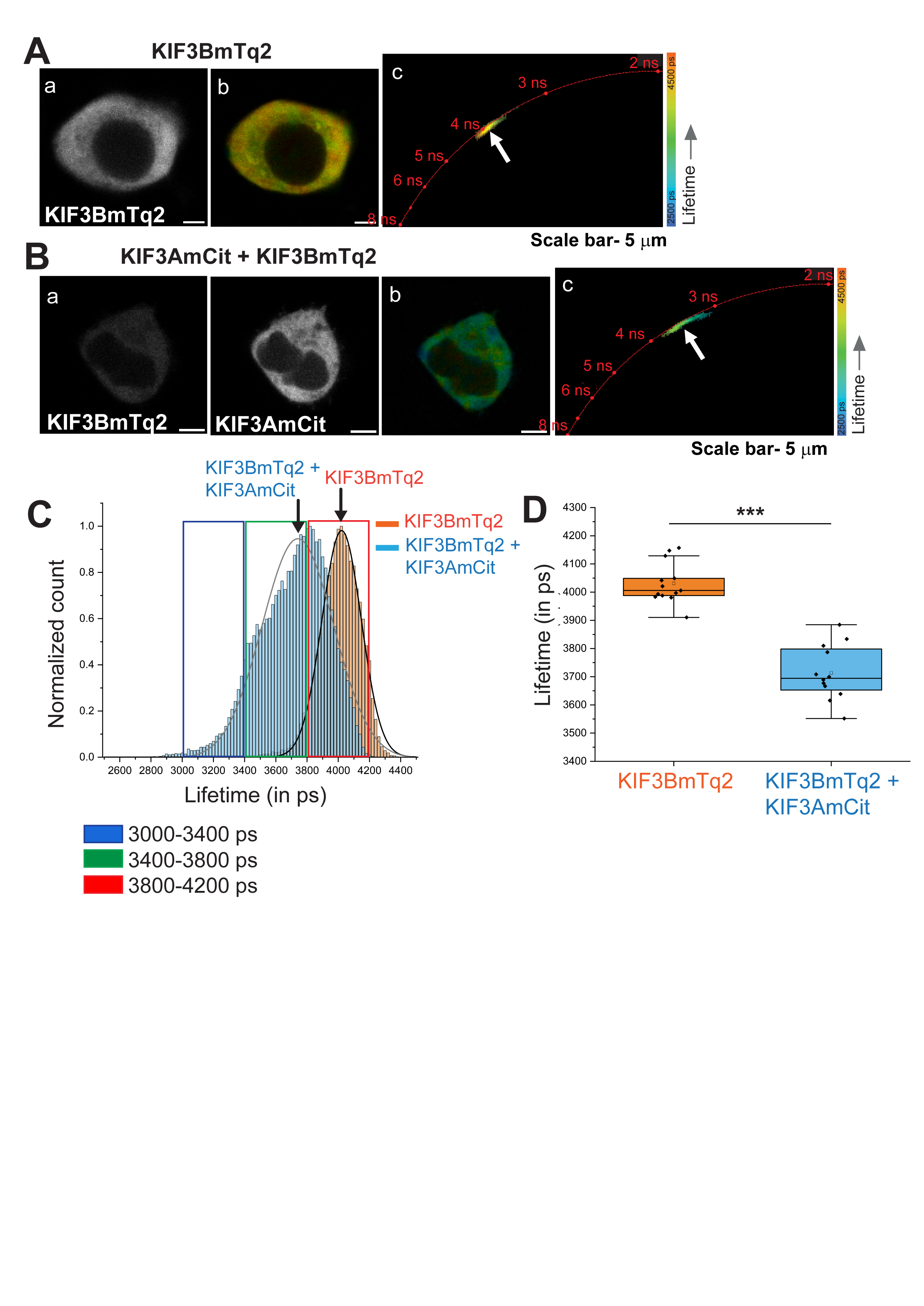

**Supplementary figure S2. Characterization of fluorescence lifetime assay to study the interaction with KIF3B using FLIM-FRET**

**A.** Fluorescence lifetime analysis of KIF3BmTq2 in the absence of an interacting partner. **(a)** Confocal image of a representative cell expressing KIF3BmTq2; **(b)** Fluorescence lifetime image of the cell where the colour depicts the fluorescence lifetime of KIF3BmTq2 as per the scale provided towards of the right of the panel; **(c)** phasor plot showing the lifetime distribution of KIF3BmTq2, arrow points to the centre of the distribution. Scale bars ~ 5 μm.

**B.** Fluorescence lifetime analysis of KIF3BmTq2 in the presence of its known interacting partner KIF3AmCit. **(a)** Confocal images of a representative cell expressing KIF3B-mTq2 and KIF3AmCit; **(b)** Fluorescence lifetime image of the cell depicting the fluorescence lifetime of KIF3BmTq2 as per the scale provided towards the right; **(c)** phasor plot showing the lifetime distribution of KIF3BmTq2, arrow points to the centre of the distribution. Scale bars ~ 5 μm.

**C.** Histogram plot showing the frequency distribution of fluorescence lifetime of KIF3BmTq2 in the absence (orange) and presence (blue) of its interacting partner KIF3AmCit.

**D.** Box plot analysis for lifetime of KIF3BmTq2 in absence (orange) (n=11 cells) and presence (blue) (n=24 cells) of KIF3AmCit. The pair-wise significance of difference was estimated using Mann-Whitney U test. *** indicates p value < 0.001.

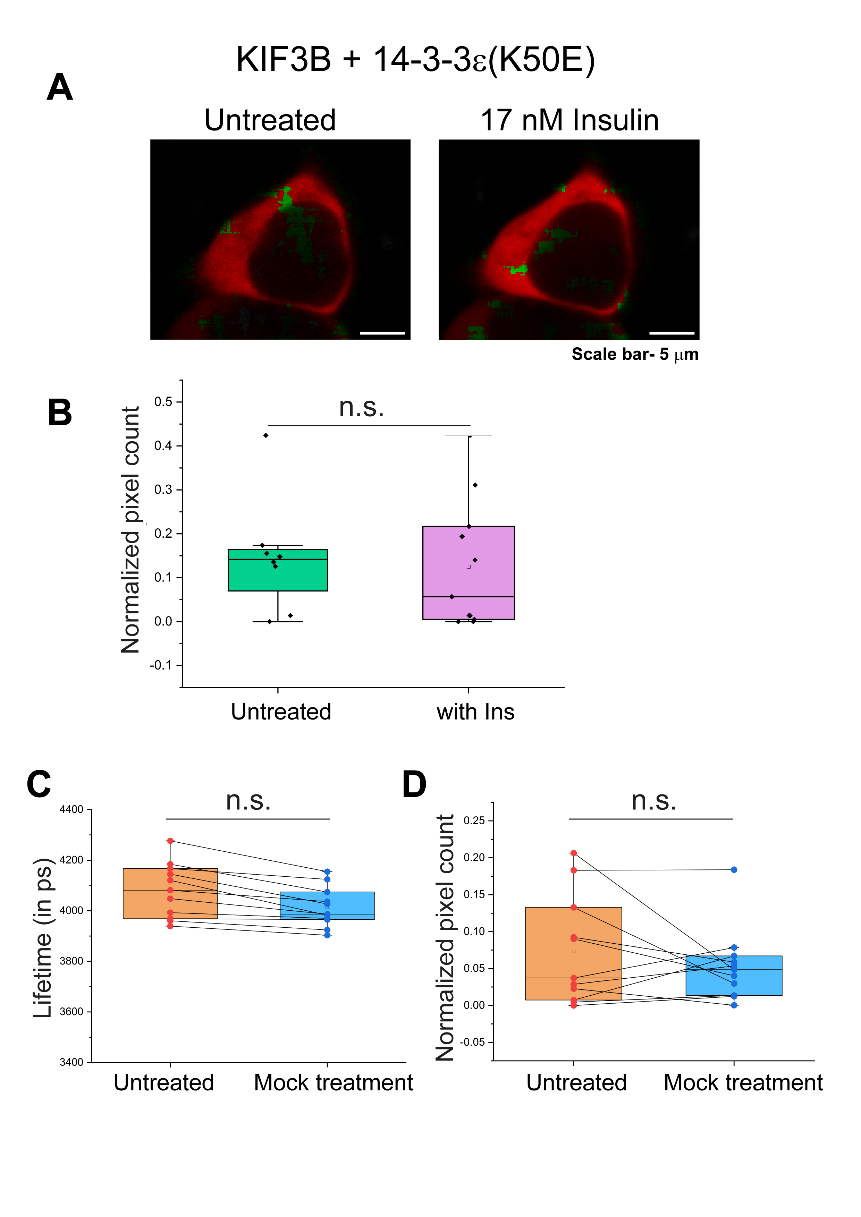

**Supplementary figure S3. Effect of insulin treatment on the lifetime of KIF3BmTq2 in the presence of 14-3-3ε(K50E)-mCit.**

**A.** The lifetime map of cells transfected with KIF3BmTq and 14-3-3ε(K50E)mCit before and after 17nm insulin treatment and shown in the RGB scale as in Figure 3.

**B.** Box and whiskers plots show the fraction of pixels with the fluorescence lifetime of KIF3BmTq2 in the range of 3000–3800 ps before and after the insulin treatment as indicated along the x-axis in the presence of 14-3-3ε(K50E)mCit. The statistical significance of difference was estimated using Mann-Whitney U test. n.s. indicates p-value > 0.05. Note that the K50E mutant is deficient in target peptide binding. Therefore, the insulin treatment failed to reduce the lifetime of KIF3BmTq2. This result is a confirmation of the insulin-induced enhancement of the interaction between KIF3B and 14-3-3ε. N=3, n=8 cells for untreated and 11 cells for with Insulin

**C.** Box and whiskers plots showing the fluorescence lifetime of KIF3BmTq2 in cells expressing both KIF3BmTq2 and 14-3-3εmCit before and after mock treatment, as indicated on the x-axis. Blank PBS was added to the cells for mock treatment.

**D.** Box and whiskers plots show the fraction of pixels with the fluorescence lifetime of KIF3BmTq2 in the range of 3000–3800 ps before and after the mock treatment, as indicated on the x-axis. N=3, n=11 cells

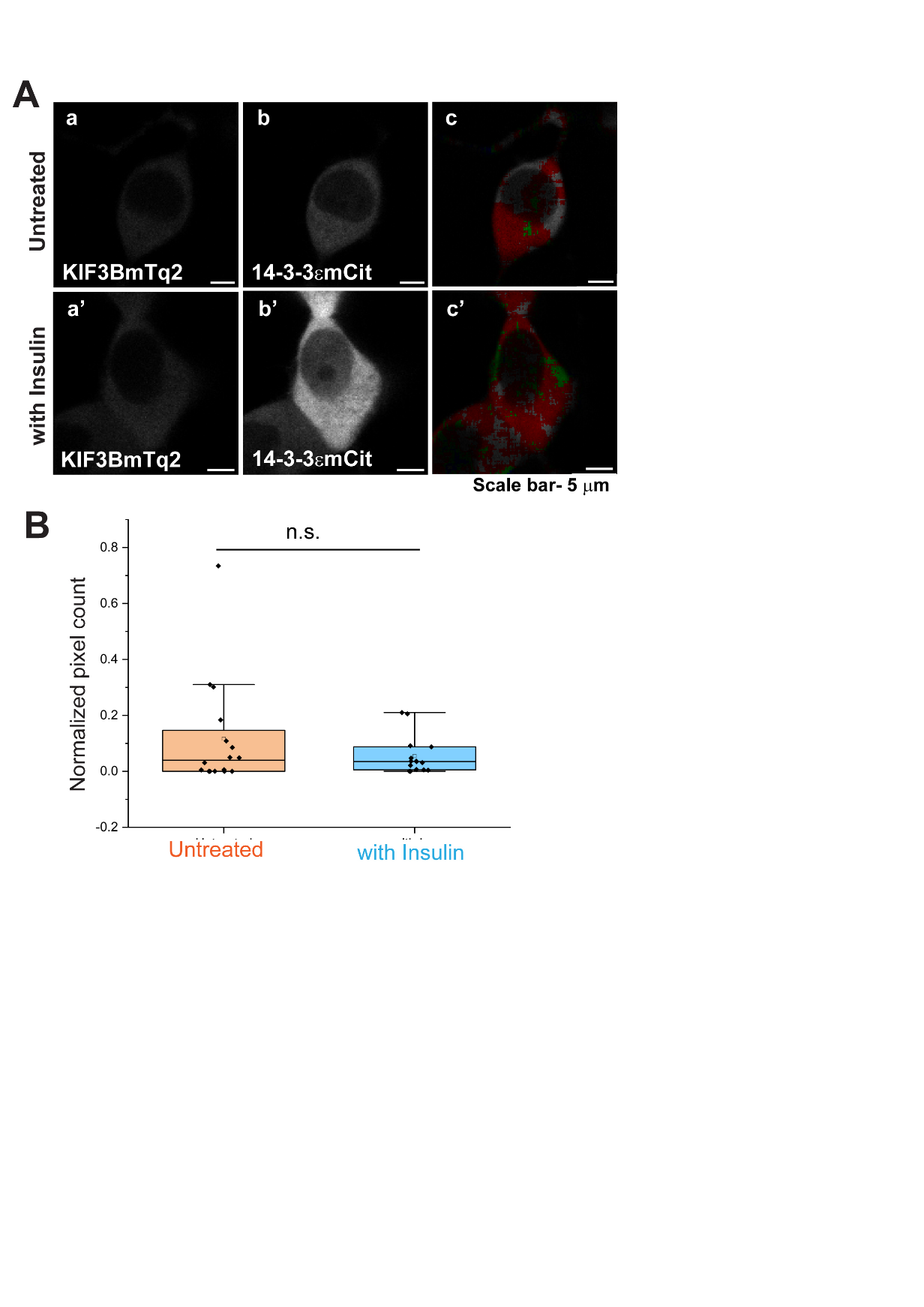

**Supplementary figure S4. Effect of insulin stimulation on the interaction between KIF3BmTq2 and 14-3-3εmCit in serum-starved cells**

**A.** Fluorescence lifetime image of KIF3BmTq2 in cells coexpressing 14-3-3εmCit before (top panel) and after (bottom panel) insulin treatment. **(a, a’)** a representative confocal image showing the expression of KIF3BmTq2, **(b, b’)** a representative confocal image showing the expression of 14-3-3εmCit. **(c, c’)** The false-colour image representing the lifetime values of KIF3BmTq2 where red represents lifetime values in range of 3800 – 4200 ps and green represents lifetime values in range of 3400 – 3800 ps. Arrow highlights the region of weak interaction between KIF3BmTq2 and 14-3-3mCit. Scale bars ~ 5 μm

**B.** Box plot analysis of lifetime of KIF3BmTq2 in untreated cells (orange) and after insulin treatment (blue). Statistical significance was tested using Unpaired t-test and Mann-Whitney U test and the data was found to be non-significant using both the tests. n.s. indicates p > 0.05. N=3, n=16 cells for untreated and 15 cells for with Insulin.

**Supplementary figure S5**

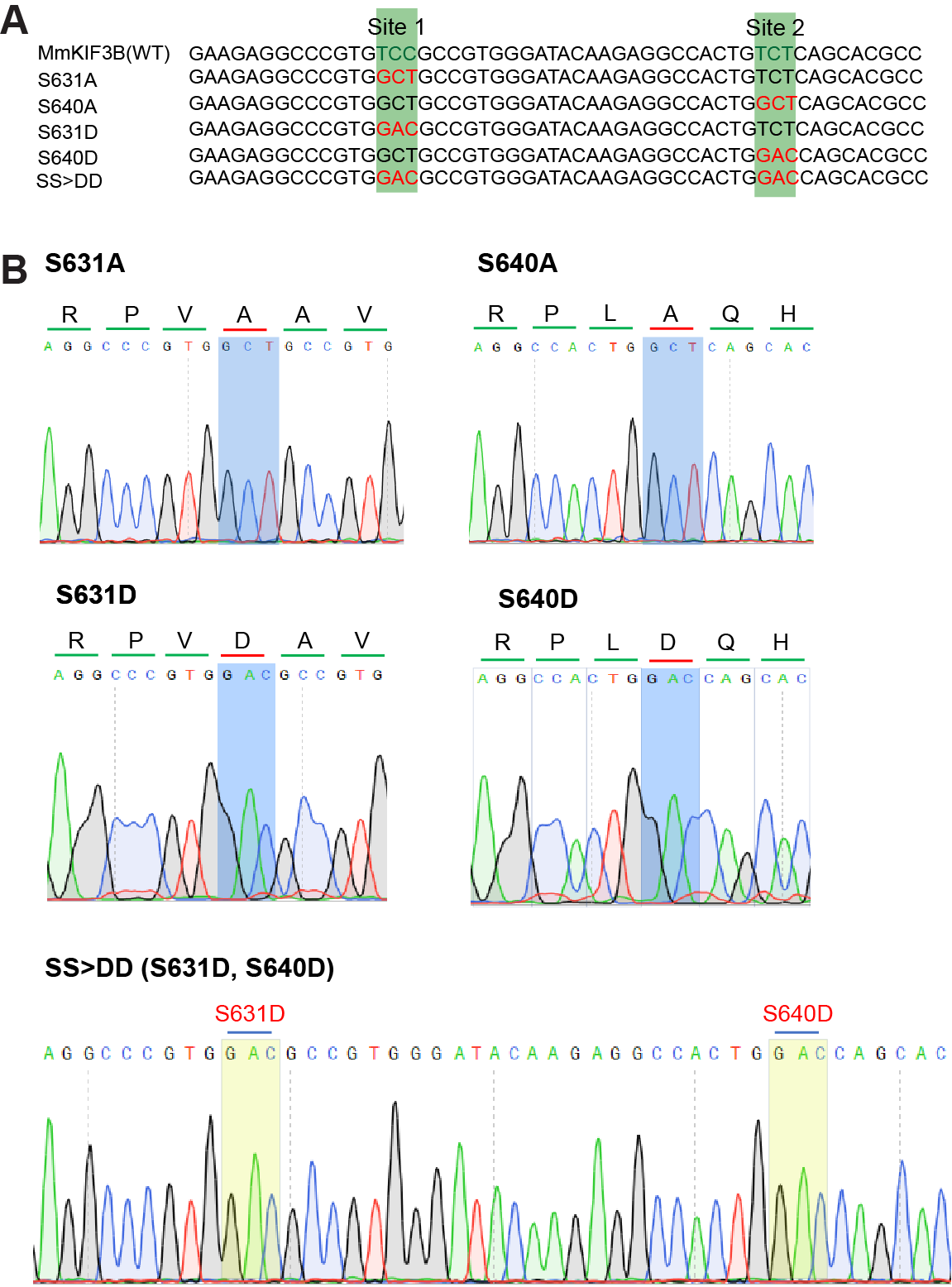

**Supplementary figure S5: Confirmation of KIF3B mutants by sequencing**

**A.** Sequence alignment of KIF3BmTq2 mutants with the wild type template p220ME_MmKIF3B(fl)-mCherry sequence, with the codons for the mutated serine residues highlighted in red.

**B.** Chromatograms show confirmation of the point mutations in the KIF3BmTq2 constructs. The codons coding for the mutated residues are marked in blue for single mutants and yellow for double mutants.

**Supplementary figure S6**

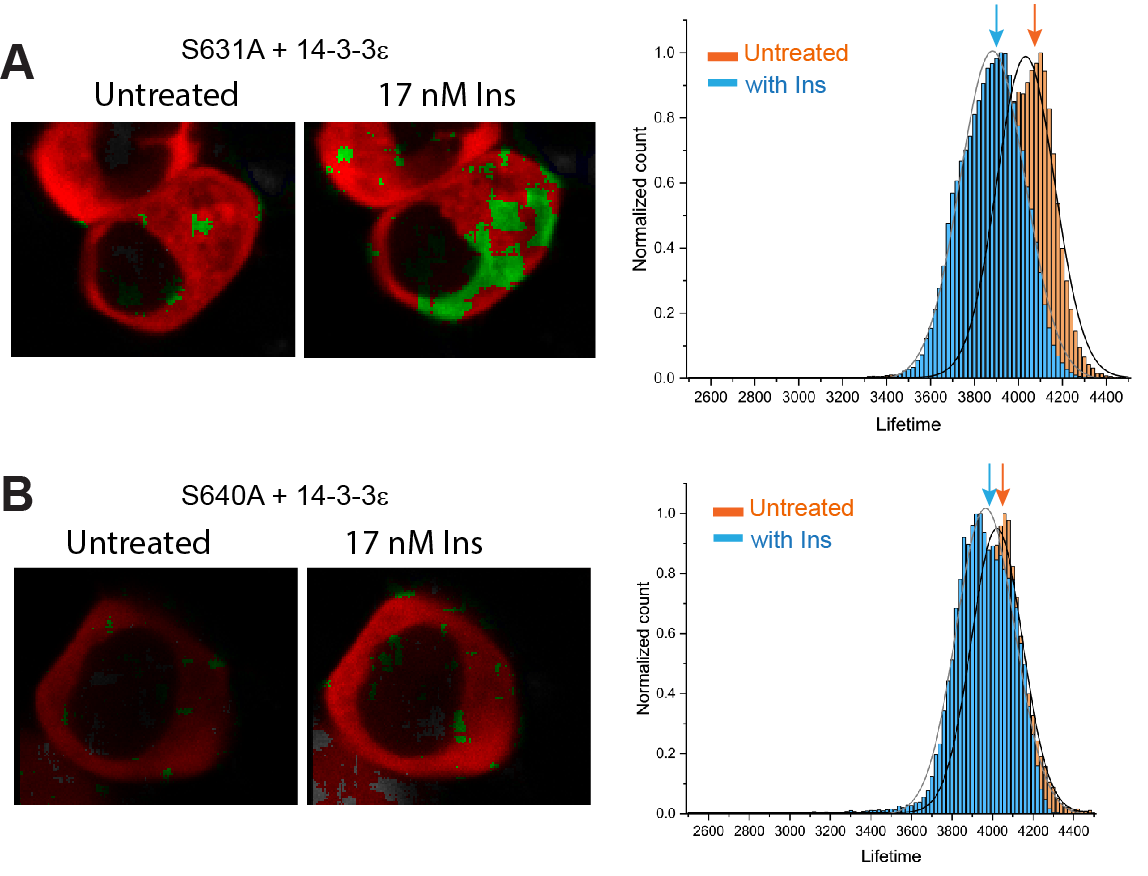

**Supplementary figure S6: Insulin treatment selectively reduces mean lifetime of KIF3Bm(631A)Tq2 in the 14-3-3εmCit background.**

**A.** Lifetime images shown in RGB scale of a cell transfected with KIF3B(S631A)mTq2 and14-3-3εmCit before and after insulin treatment. The histogram distribution plot adjoining the images show shift in the mean lifetime values obtained from each pixel from n=34 cells for untreated and 32 cells for with Insulin (N=3).

**B.** Lifetime images shown in RGB scale of a cell transfected with KIF3B(S640A)mTq2 and14-3-3εmCit before and after insulin treatment. The histogram distribution plot adjoining the images shows the frequency distribution of lifetime from each pixel obtained from n=13 cells for untreated and 14 cells for with Insulin (N=3).

**Supplementary figure S7**

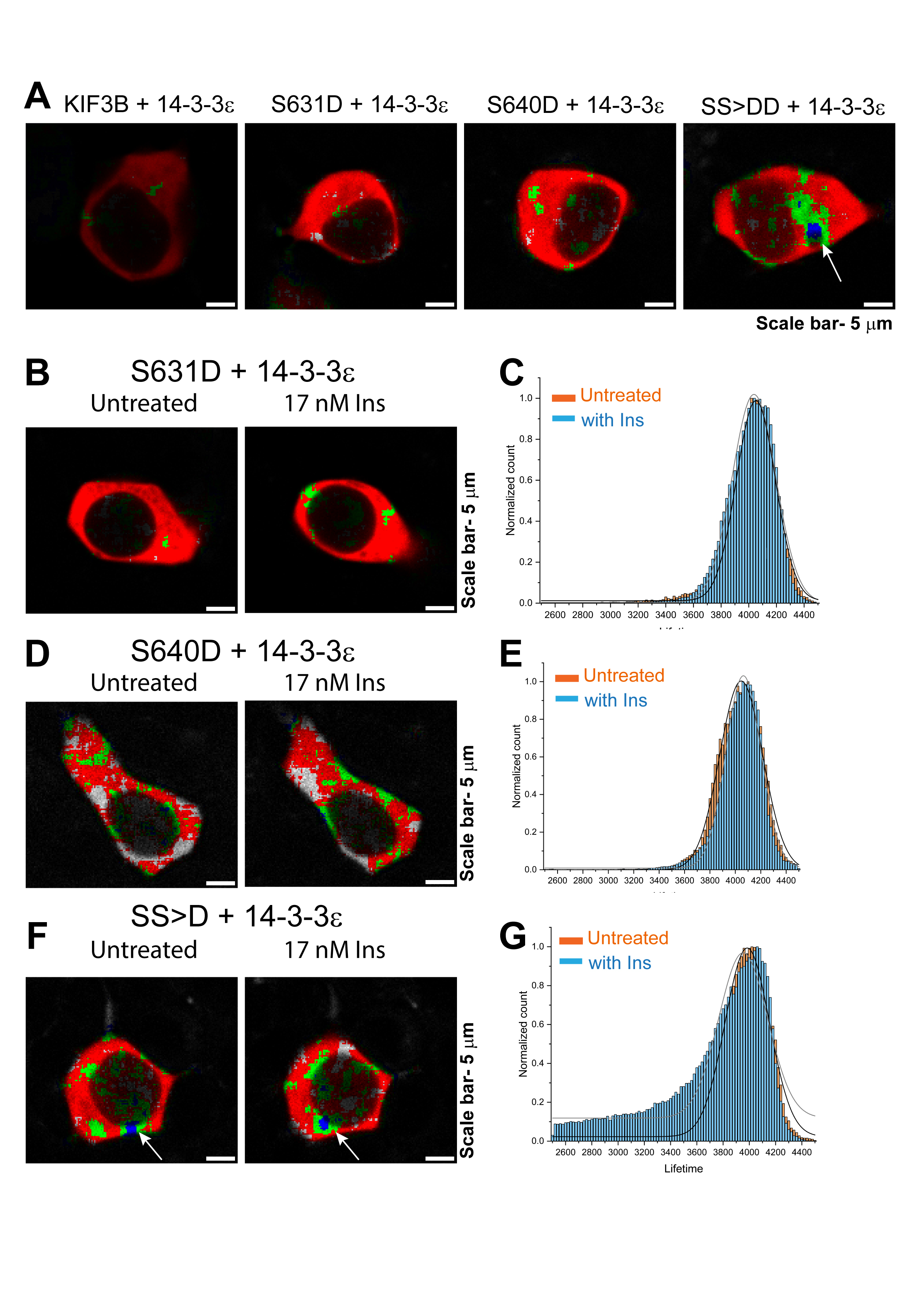

**Supplementary figure S7: Insulin treatment did not reduces mean lifetime of KIF3BmTq2 aspartate substitution mutants in the 14-3-3ε-mCit background.**

A. Lifetime image shown in RGB colour scheme of representative cells co-transfected with wild type and S631D, S640D and SS>DD aspartate mutants of KIF3B with 14-3-3εmCit, respectively, as indicated on the top of each image. Scale bar ~ 5 μm. N=3

**B, D, F.** Representative images showing the lifetime distribution in the cells co-transfected with KIF3B(S631D)mTq2, KIF3B(S640D)mTq2 and KIF3B(SS>DD)mTq2 with 14-3-3εmCit, respectively, before and after insulin treatment, as indicated on top of each image. Scale bar ~ 5 μm

**C, E, G.** Histogram plot showing the frequency distribution of lifetime from each pixel of cells co-transfected with the plasmid combination in the adjoining images, before (orange bars) and after insulin treatment (blue bars) obtained from multiple cells (N ≥ 2). KIF3B(S631D)mTq2+14-3-3εmCit: untreated- 10 cells, with Insulin- 18 cells; KIF3B(S640D)mTq2+14-3-3εmCit: untreated- 11 cells, with Insulin- 21 cells; KIF3B(SS>DD)mTq2+14-3-3εmCit: untreated- 9 cells, with Insulin- 17 cells

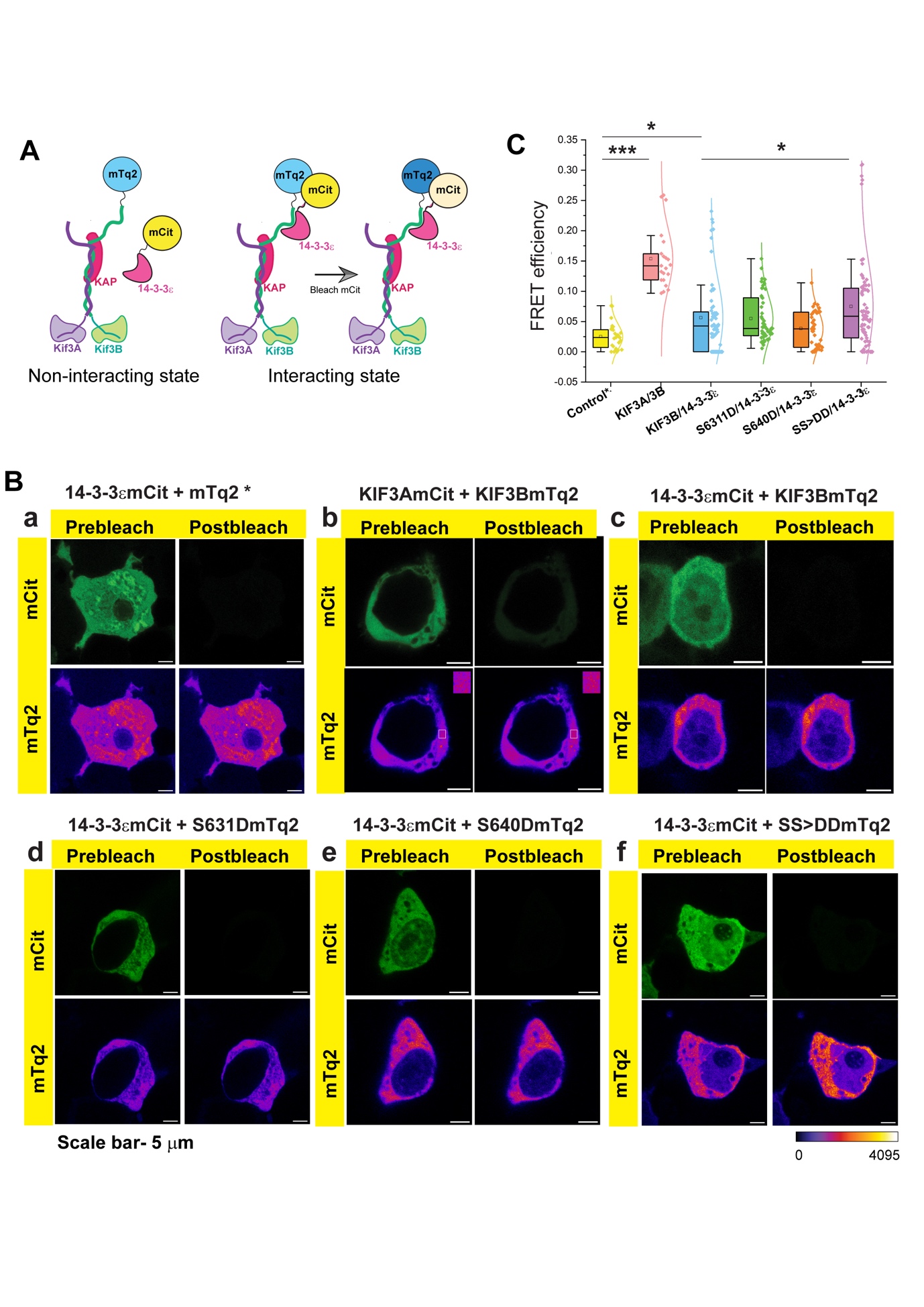

**Supplementary figure S8. Acceptor photobleach (AP)-FRET analysis shows that phosphorylation at Ser631 and Ser640 play a role in interaction of 14-3-3**ε **with KIF3B.**

**A.** Schematic representation of AP-FRET assay where KIF3B is tagged with mTurquoise2 and 14-3-3ε is tagged with mCitrine.

**B.** The image panels show the prebleach and postbleach intensity of mTurquoise2 (Fire LUT) and mCitrine (green) in different combinations of transfections as indicated on the top of each group panel. The mTq2 and 14-3-3εmCit **(a)** was used as negative control, and the pair of KIF3BmTq2 with KIF3AmCit **(b)** was used as the positive control. The insets in **(b)** show a magnified view of the region marked by the white box in both prebleach and postbleach mTurquoise2 channel respectively, to highlight the increase in postbleach intensity. The wild type or aspartate mutants of KIF3BmTq2 were co-transfected with 14-3-3εmCit in different combinations as shown on top of each panel **c-f**, and AP-FRET assay was performed in cells expressing both the fluorophores. Scale bars ~ 5 µm.

**C.** The box and whisker plots and associated distribution fit on the actual measurements depict the quantitative differences in the FRET efficiencies in each condition as depicted in panel C (a-f) on a relative scale indicated on the y-axis. Each dot represents the FRET efficiency calculated in a randomly selected ROI in the cytoplasm of a selected cell. ROIs from multiple cells were taken to generate the data. The pair-wise significance of difference was estimated using Mann-Whitney U test. * Indicates p value < 0.05 and *** indicates p value < 0.001. The central line shows the median. The whiskers show the values extending to 1.5X the interquartile range from the Q1 or Q3 boundaries. n>30 ROIs for each case.

**Supplementary figure 9**

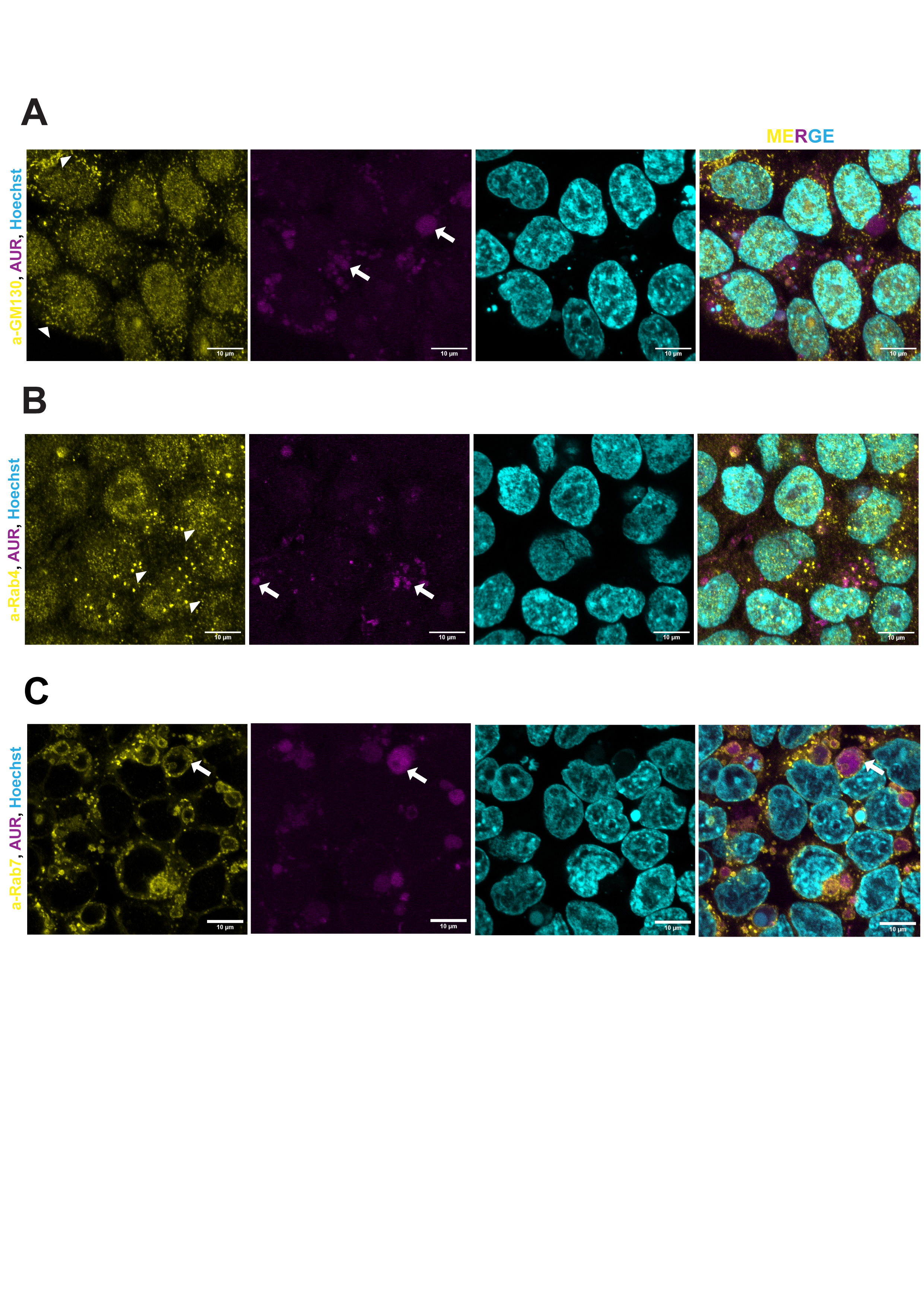

**Supplementary figure S9: Immunostaining coupled with AUR assay in HEK293T cells reveals the compartment of interaction between KIF3B and 14-3-3**ε **are Rab-7 positive**

**A-C.** Images show the result for antibody staining (in yellow) and AUR assay (in magenta) in HEK293T cells co-transfected with 14-3-3ε-S1 and KIF3B-S2. The nuclei are stained with Hoechst and shown in cyan. α-GM130, α-Rab-4 and α-Rab7 were used for immunostaining in panel A, B and C, respectively. Note that the resorufin signal in panel **C** is encircled by Rab7 staining, suggesting KIF3B and 14-3-3ε might interact with each in Rab7-positive compartments

**Table S1 :** **List of constructs used in the study, made by subcloning**

| **S. no.** | **Plasmid name** | **Insert name** | **Source of Insert** | **Backbone/Destination vector** | **Restriction sites** |
| --- | --- | --- | --- | --- | --- |
| 1. | KIF3BmTq2 | mTurquoise2 | mKOκ-2A-mTurquoise2 | p220ME_MmKIF3B(fl)-mCherry | BamH1, Not1 |
| 2. | KIF3AmCitrine | Full length KIF3A | p285ME_MmKIF3A(fl) | mCitrine-N1 | Nhe1, Age1 |
| 3. | 14-3-3εmCitrine | 14-3-3ε | GST-14-3-3ε | mCitrine-N1 | Xho1, Kpn1 |
| 4. | 14-3-3ε-K50E-mCitrine | 14-3-3ε-K50E | GST-14-3-3ε-K50E | 14-3-3εmCitrine | Xho1, Kpn1 |
| 5. | mTurquoise2 | mTurquoise2 | KIF3BmTq2 | KIF3BmTq2 | Nhe1, Not1 |
| 6. | 14-3-3ε-S1 | i) 14-3-3ε  ii) S1 | i) 14-3-3εmCitrine  ii) mito-V5-APEX2 | KIF3BmTq2 | i) Nhe1, Xho1  ii) Xho1, Not1 |
| 7. | KIF3A-S1 | i) KIF3A  ii) S1 | i) p285ME_MmKIF3A(fl)  ii) mito-V5-APEX2 | pIRES | i) Xho1, Mlu1  ii) Mlu1, Xba1 |
| 8. | KIF3B-S2 | S2 | mito-V5-APEX2 | p220ME_MmKIF3B(fl)-mCherry | BamH1, Not1 |

**Table S2 :** **List of primers used for site-directed mutagenesis**

| **S. no.** | **Plasmid name** | **Forward primer for SDM** | **Reverse primer for SDM** |
| --- | --- | --- | --- |
| 1. | KIF3Bm(S631A)Tq2 | CCGTG**GCT**GCCGTGGGATACAAGAGG | CACGGC**AGC**CACGGGCCTCTTCATCATCTGCTGG |
| 2. | KIF3B(S640A)mTq2 | GCCACTG**GCT**CAGCACGCCAGAATGAGCATG | GTGCTG**AGC**CAGTGGCCTCTTGTATCCCACGGC |
| 3. | KIF3B(S631D)mTq2 | CCGTG**GAC**GCCGTGGGATACAAGAGG | CACGGC**GTC**CACGGGCCTCTTCATCATCTGC |
| 4. | KIF3B(S640D)mTq2 | GCCACTG**GAC**CAGCACGCCAGAATGAGCATG | GTGCTG**GTC**CAGTGGCCTCTTGTATCCCACGGC |
| 5. | 14-3-3ε-K50E-S1 | GCATAT**GAA**AATGTGATTGGAGCTAGAAGAGCCTCC | CACATT**TTC**ATATGCAACAGATAGGAGGTTTCTTTC |

**Table 3: List of other reagents used in the study**

| **S. no.** | **Reagent name** | **Company** | **Catalogue number** |
| --- | --- | --- | --- |
| 1. | DMEM | Gibco-Thermo Fisher Scientific Inc., USA | **11965-092** |
| 2. | FBS | Gibco-Thermo Fisher Scientific Inc., USA | **1600044** |
| 3. | PenStrep | Gibco-Thermo Fisher Scientific Inc., USA | **15140** |
| 4. | Lipofectamine 3000 | Thermo Fisher Scientific Inc. | **L3000015** |
| 5. | Amplex UltraRed | Thermo Fisher Scientific Inc. USA | **A36006** |
| 6. | Vectashield | Vector laboratories | **H-1000** |
| 7. | Insulin | Sigma-Aldrich Chemicals Pvt. Ltd | **I9278** |
| 8. | Hoechst | Thermo Fisher Scientific  Inc.,USA | **33342** |
| **9.** | α-14-3-3ε | Chemicon | **AB9734** |
| **10.** | α-GST | Genei | **M0101** |
| **11.** | α-GM130 | Thermo Fisher Scientific | **PA-077** |
| **12.** | α-Rab4 | Thermo Fisher Scientific | **PA3-912** |
| **13.** | α-Rab7 | Abclonal | **ab12308** |
| **14.** | α-Rabbit-HRP | Genei | **HP020** |
| **15.** | α-Mouse-HRP | Genei | **HP06** |
| **16.** | α-Rabbit-Oregon Green 488 | Life technologies | **011038** |
| **17.** | Normal Rabbit IgG | Cell Signaling Technology | **2729S** |
| **18.** | α-KIF3A | Cell Signaling Technology | **8507S** |
| **19.** | α-14-3-3ε | Santa Cruz Biotechnology | **sc- 23957** |
| **20.** | α-KIF3B | Santa Cruz Biotechnology | **sc- 514165** |
| **21.** | α-6X-His | Abcam | **ab 18184** |
